# Tracing the origin of Oriental beech stands across Western Europe and reporting hybridization with European beech – implications for assisted gene flow

**DOI:** 10.1101/2022.07.25.501368

**Authors:** Mirjam Kurz, Adrian Kölz, Jonas Gorges, Beatriz Pablo Carmona, Peter Brang, Yann Vitasse, Martin Kohler, Fabio Rezzonico, Theo H. M. Smits, Jürgen Bauhus, Andreas Rudow, Ole Kim Hansen, Mohammad Vatanparast, Hakan Sevik, Petar Zhelev, Dušan Gömöry, Ladislav Paule, Christoph Sperisen, Katalin Csilléry

## Abstract

Human-aided translocation of individuals within the species’ range, assisted gene flow (AGF), has been suggested as a climate change mitigation strategy, especially for foundational species, such as forest trees. The benefits and risks of AGF largely depend on the genetic divergence between host and donor populations, their rate and direction of hybridization, and the climate distance that the transfer involves. In this study, we explored the use of Oriental beech (*Fagus sylvatica* subsp. *orientalis*), growing from Iran to the Balkans, for AGF in populations of European beech (*F. sylvatica* subsp. *sylvatica*), which grow throughout Europe and are increasingly affected by climate warming. Using 16 microsatellite loci and samples from 13 and 6 natural populations of Oriental and European beech, respectively, we identified 5 distinct genetic clusters in Oriental beech with a divergence (F_ST_) of 0.15 to 0.25 from European beech. Using this knowledge, we tracked the origin of 11 Oriental beech stands in Western Europe, some established in the early 1900s. In two stands of Greater Caucasus origin, we additionally genotyped offspring and found evidence for extensive hybridization, with 41.3% and 17.8% of the offspring having a hybrid status. Further, climate data revealed a higher degree of seasonality across the Oriental beech growing sites than across the planting sites in Western Europe, with some sites additionally having a warmer and drier climate. Accordingly, in one of these stands, we found evidence that bud burst of Oriental beech occurs four days earlier than in European beech. These results suggest that AGF of Oriental beech could increase the genetic diversity of European beech stands and may even help the introgression of variants that are more adapted to future climatic conditions. Our study showcases an evaluation of the benefits and risks of AGF and calls for similar studies on other native tree species.

## Introduction

Rapid climate change has already compromised the persistence of many tree species in their current distributional range and this trend is expected to increase in the future (Masson-Delmotte et al., 2021). In Europe, for example, drought caused approximately 500,000 ha of excess forest mortality between 1987 and 2016 (Senf et al., 2020), with long-term consequences for ecosystem services (Hanewinkel et al., 2013; Morin et al., 2018; Seidl et al., 2017). While range expansion to higher elevations has been observed for many forest tree species (Vitasse et al., 2021), their migration towards higher latitudes requires a much larger distance to track a similar thermal change, which is often hampered by human land use and habitat fragmentation (Miller & McGill, 2018). It is therefore widely agreed that human interventions are necessary to maintain healthy and productive forests (Brang et al., 2014; Linder, 2000).

Assisted migration (AM) is the human-aided translocation of individuals to mitigate the adverse effects of climate change or resistance to diseases (McLachlan et al., 2007; Peters, 1985). Established conservation paradigms favor maintaining the status quo of perceived natural ranges of species and prefer in situ conservation and management. Thus, AM has been refused by many conservation experts for reasons related to the lack of knowledge about long-term consequences of introducing exotic materials, to costs, and to ethical issues (Hagerman et al., 2010; Ricciardi & Simberloff, 2009). Introductions of non-native tree species, often from a different continent with no local closely related species, are the oldest examples of AM. In these situations, the expected benefits are primarily related to a few desired ecosystem services and the risks are largely ecological, such as disease and pest outbreaks due to novel host–pest interactions or the introduced species becoming invasive (Winder et al., 2011).

More recently, due to the rapid expansion of genetic data sets to study adaptation (Hoffmann & Willi, 2008; Savolainen et al., 2013) and the pressure from unprecedented climate change, the concept and evaluation of AM has evolved. In particular, a type of AM, assisted gene flow (AGF), where individuals are moved within the species’ range, is increasingly put forward as a potential management tool to mitigate maladaptation (Aitken & Whitlock, 2013; Hamilton & Miller, 2016), especially for long-living organisms such as forest trees that are unable to track their climatic niche (Aitken & Bemmels, 2016). The main benefits and risks of AGF are related to the genetic composition of host and donor populations (Aitken & Whitlock, 2013; Weeks et al., 2011). The benefits can be either increased genetic diversity, and therefore also higher adaptive potential (Sgrò et al., 2011), and/or the introduction of variants preadapted to future expected conditions. Despite overwhelming empirical evidence for negative fitness and compromised disease resistance of populations with reduced genetic diversity (Aerts & Honnay, 2011; Smulders et al., 2008; Zeng & Fischer, 2021), many afforestation and forest restoration programs still incorporate only a few seed families (Bettinger et al., 2009; Sperisen et al., 2016). Thus, practitioners might be especially resistant to the introduction of different provenances to increase genetic diversity. Introducing preadapted genetic variants has long been proposed in forestry, using transfer functions that relate trait observations from provenance trials to the climatic distance of the transferred provenance (e.g., Mátyás 1994) or, more recently, using genomic data (e.g., Mahony et al., 2020). However, thus far, no large-scale introductions of foreign provenances have been performed in Europe (Gömöry et al., 2021) or North America (e.g., Pedlar et al., 2020).

The choice of the introduced provenance is a key factor in any AGF program because its benefits, as well as its risks, are expected to increase with the genetic divergence between host and donor populations. After a certain degree of divergence, such as with subspecies, we start referring to hybridization and introgression between the host and donor gene pools (Abbott et al., 2013; Barton, 2008). Hybridization increases genetic diversity, enables the introgression of preadapted alleles into a population, and may create beneficial gene interactions; it is thus considered a major source of adaptation (Aitken & Bemmels, 2016; Broadhurst et al., 2008; Kremer & Hipp, 2020; Tigano & Friesen, 2016; Weeks et al., 2011). However, hybridization can also break up beneficial allele combinations, increase the risk of outbreeding depression, and lead to a reduction in offspring fitness compared with the parental generation (Edmands, 2006; Frankham et al., 2011), which can in turn accelerate the extinction of lineages or even species (Rhymer & Simberloff, 1996; Todesco et al., 2016).

Oriental beech (*Fagus sylvatica* subsp. *orientalis* (Lipsky) Greut. & Burd) and European beech (*Fagus sylvatica* subsp. *sylvatica* L.) are sister subspecies (Denk, 1999a, 1999b; Denk et al., 2002) whose combined ranges extend almost continuously from Iran to Spain (Fig. 1a). The range of European beech is fairly continuous over large parts of Europe, while Oriental beech grows in several loosely connected or completely isolated populations from the southeastern Balkans, where it is in contact with European beech, to Asia Minor and the Caucasus, to Northern Iran (Fig. 1a; Kandemir and Kaya, 2009). Morphological studies have pointed out that Oriental beech harbors higher trait diversity, in terms of leaf and cupule morphology, than European beech (Denk, 1999a; Denk et al., 2002). More recently, molecular analyses using isozymes and chloroplast DNA have shown that Oriental beech also has a higher genetic diversity (Bijarpasi et al., 2020; Denk et al., 2002b; Gömöry et al., 1999; Paffetti et al., 2007; Vettori et al., 2004). Further, isozyme data combined with fossil evidence suggests that European beech diverged from a single lineage of Oriental beech from Asia Minor, some 817 ky ago, and colonized Europe only at the beginning of the last glacial period (Gömöry et al., 2018; Magri et al., 2006).

**Figure 1.**
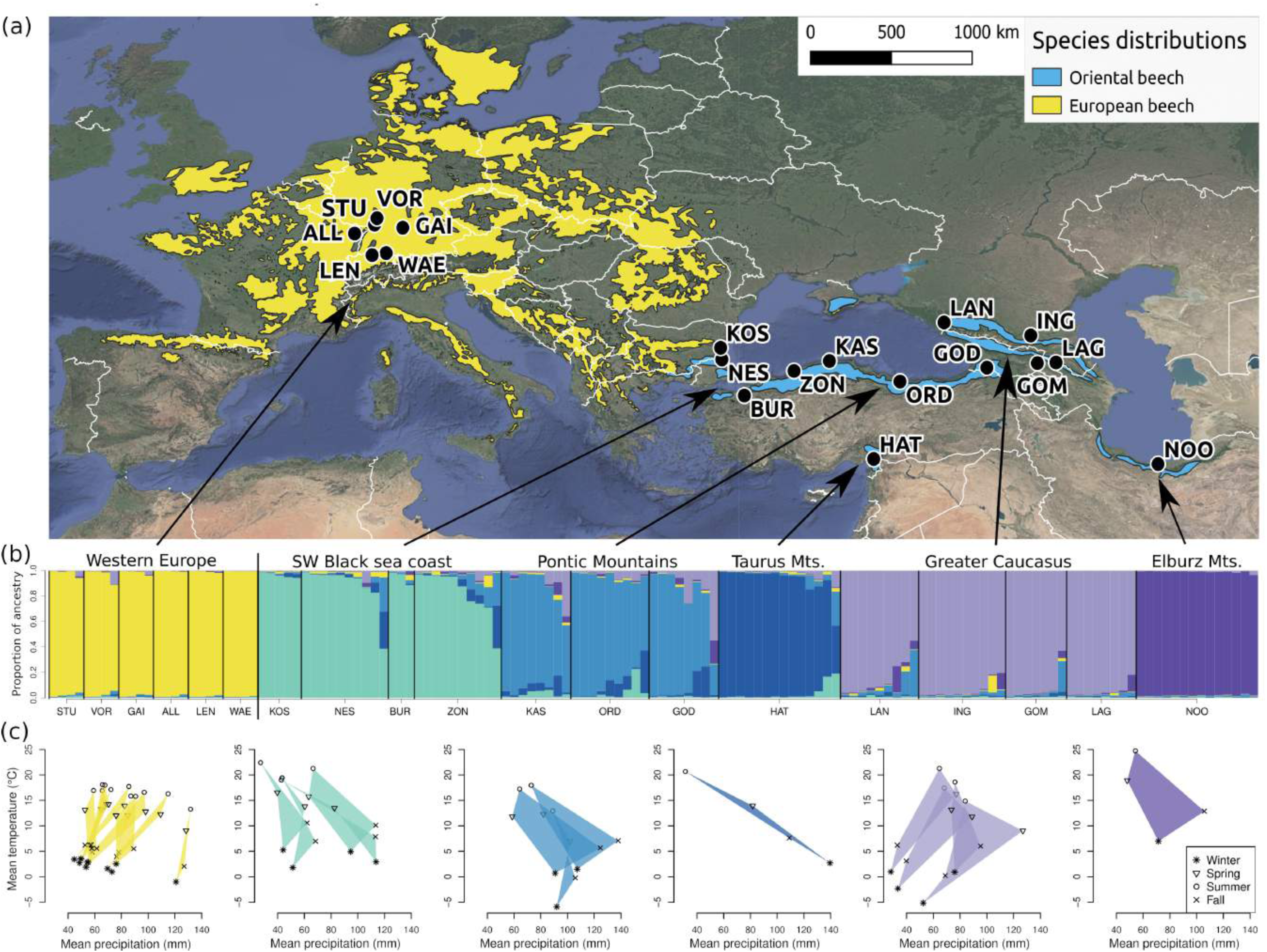
Spatial **(a)**, genetic **(b)** and climatic **(c)** characterization of the natural European and Oriental beech populations. **(a)** Overview of sampling locations and the species distribution ranges (source: EUFORGEN). **(b)** Genetic clustering of natural European and Oriental beech populations using the software Structure assuming six clusters. Each bar corresponds to a sampled individual and each color to a genetic cluster that correspond to the mountain ranges/regions listed in Table 1. **(c)** Climate of the growing sites in terms of temperature and precipitation. Polygons show the monthly mean temperature and precipitation averaged for the 3 months of each season at the sites (see Table S2 and Materials and Methods for details).

**Table 1.** Natural Oriental beech populations considered in this investigation: their locations and the abbreviations (ID) used in the study, and the number of individuals sampled from each population for genetic analyses (N). Populations are listed from east to west and from south to north.

| Mountain range<br>Region | Country | Nearby town<br>Locality | ID | Longitude | Latitude | N |
| --- | --- | --- | --- | --- | --- | --- |
| Elburz Mountains | Iran | Noor | NOO | 52°01'40.8"E | 36°34'30.3"N | 14 |
| Greater Caucasus | Georgia | Lagodechi | LAG | 46°21'25.6"E | 41°53'44.2"N | 8 |
| Greater Caucasus | Georgia | Gombori Pass | GOM | 45°19'23.7"E | 41°51'40.7"N | 7 |
| Greater Caucasus | Russia | Inguschetien | ING | 44°57'16.2"E | 43°18'44.7"N | 10 |
| Greater Caucasus | Russia | Lago-Naki | LAN | 40°08'04.2"E | 43°59'30.8"N | 9 |
| Taurus Mountains | Turkey | Dörtyol (Hatay) | HAT | 36°13'17.6"E | 36°50'36.1"N | 14 |
| Pontic Mountains | Georgia | Goderdzi Pass | GOD | 42°30'56.2"E | 41°37'35.6"N | 8 |
| Pontic Mountains | Turkey | Ordu | ORD | 37° 41' 11" E | 40° 53' 51" N | 9 |
| Pontic Mountains | Turkey | Kastamonu | KAS | 33°45'33.7"E | 41°58'38.1"N | 8 |
| SW Black Sea coast | Turkey | Zonguldak | ZON | 31°47'21.8"E | 41°27'12.7"N | 10 |
| SW Black Sea coast | Turkey | Bursa | BUR | 29°01'37.7"E | 40°09'43.5"N | 3 |
| SW Black Sea coast | Bulgaria | Kosti (Strandzha Mts.) | KOS | 27°46'51.2"E | 42°03'22.9"N | 10 |
| SW Black Sea coast | Bulgaria | Nessebar | NES | 27°42'40.1"E | 42°39'57.5"N | 5 |

To mitigate the impacts of climate change in European forests, Oriental beech has been proposed for growth trials (Schmiedinger et al., 2009; Vítková et al., 2016) and also for small-scale introductions to mitigate drought-induced forest mortality in European beech (Brang et al., 2016). Indeed, Oriental beech fills a similar ecological niche and forms similar plant associations in its native range as European beech (Willner et al., 2017), and most of the potential pathogens or pest species seem to occur equally on both beech subspecies (Felbermeier & Marvie-Mohadjer, 2010). Starting at the beginning of the 20th century, a few Oriental beech plantations and beech provenance trials with Oriental beech were established in Europe and suggested that Oriental beech may exhibit growth superior to European beech when facing drought periods (Bogunović et al., 2020; Elzami, 2018; Gorges, 2020). Thus, the two species appear to support a very similar flora, fauna and fungi, and hence the ecological risks of the introduction of Oriental beech are likely small. In contrast, the genetic benefits and risks could be important and likely depend on the provenance of Oriental beech. Indeed, Oriental beech is best regarded as a species complex, and different populations are expected to have a different degree of divergence from European beech (Gömöry & Paule, 2010). Thus, the provenance of the introduced Oriental beech should play a pivotal role in determining the potential gain in genetic diversity and adaptability versus the risk of outbreeding depression due to hybridization between the host and donor populations.

The overall aim of this study was to provide a basis for the evaluation of benefits and risks of AGF for Oriental beech in Western and Central Europe, using existing forest stands as natural laboratories. Our specific aims were to: (1) expand our knowledge about the genetic diversity and structure of Oriental beech across its range using microsatellite loci; (2) establish a database of origin-tracked planted Oriental beech stands in Western Europe, complemented with knowledge about the degree of divergence between native European and introduced Oriental beech and about the climate at the planting and donor sites; (3) assess the extent to which the two subspecies hybridize upon introduction of Oriental beech in two of the oldest stands established in the early 1900s; and (4) assess phenological differences between the two subspecies to better anticipate the direction in which hybridization may occur. Concerning the last two aims, we hypothesized that European and Oriental beech would hybridize because the two subspecies are known to hybridize in their natural contact zone in the Southeastern Balkans (Cardoni et al., 2021; Müller et al., 2019). Further, since European beech starts to reproduce at the age of 40–50 years (Houston Durrant et al., 2016), we expected that we would principally find F1 hybrids, but also possibly backcrosses given the age of 100 years of the studied stands.

## Materials and Methods

### Sampling

We sampled leaves from 13 natural Oriental beech populations (Fig. 1, Table 1). Our sampling sites for Oriental beech were selected to capture genetic variation across the entire species range, and were situated in the Elburz Mountains in northern Iran, Greater Caucasus in Russia and Georgia, eastern Mediterranean region of Turkey (Taurus Mountains), southern Black Sea coast with the Pontic Mountains from Georgia to northern Turkey, and the western Black Sea coast of Turkey and Bulgaria. We successfully genotyped between 3 and 14 individuals from each Oriental beech population (Table 1).

We created a network composed of 11 planted Oriental beech stands with mature, seed-producing trees located in Switzerland, France and Germany (Table 2). We extracted information on Oriental beech stands in Switzerland from the Swiss database for exotic tree species (Bürgi & Diez, 1986). One additional stand in Switzerland (Wäldi; WAE) was reported to us by the cantonal forest service (Ulrich Ulmer, personal communication). In France, we identified one stand (Allenwiller; ALL) in northern Alsace (Klein, 1981). In Germany, we identified eight Oriental beech stands with the help of individual contacts in the state forest services and at forest research stations. In most of the identified stands, Oriental beech was admixed with European beech. At three sites (WUP, RIE, GRA), Oriental beech was growing in pure stands adjacent to European beech stands (Table 2).

**Table 2.** Oriental beech plantations in Switzerland (CH), France (FR) and Germany (D): their location; characteristics of the forest stand, including the size of the area with Oriental beech, type, age and management; and the total number of adult trees sampled for genetic analyses (N). Note that N includes both European and Oriental beech. Most plantations are mixed, meaning that Oriental and European beech trees are mixed in space, but the degree of mixing varies across plantations. In pure stands, all Oriental beech trees are strictly limited to one area without the presence of European beech. Plantations are listed in alphabetical order for each country.

| Country | Nearby village | ID | Lat | Lon | Elevation (m a.s.l.) | Stand size (ha) | Stand type | Year of planting | Management history | N |
| --- | --- | --- | --- | --- | --- | --- | --- | --- | --- | --- |
| CH | Lengnau | LEN | 47°12'00.0"N | 7°20'24.0"E | 415 | 0.2 | mixed | approx. 1920 | Final cutting due to storm damage | 18 |
| CH | Wäldi | WAE | 47°37'46.0"N | 9°06'17.4"E | 610 | 0.1 | mixed | 1921 | Even aged | 12 |
| FR | Allenwiller | ALL | 48°39'00.0"N | 7°21'00.0"E | 314 | 0.4 | mixed | 1923 | Thinning every 10 years | 16 |
| D | Gaishardt | GAI | 48°57'36.0"N | 10°01'48.0"E | 550 | 1.7 | mixed | approx. 1920 | Thinning every 10 years | 30 |
| D | Grafenberg | GRA | 48°33'34.13"N | 9°19'5.52"E | 419 | 0.1 | pure | approx. 1990 | Initial tending | 10 |
| D | Leiselheim | LEI | 48°07'12.0"N | 7°38'24.0"E | 257 | 0.5 | mixed | approx. 1920 | Thinning every 10 years | 12 |
| D | Kirkel | KIR | 49°28'47.02"N | 7°26'35.21"E | 866 | 0.9 | mixed | 1900 | A biosphere reserve with no interventions since 2009 | 10 |
| D | Ried | RIE | 50°01'12.0"N | 8°43'48.0"E | 180 | 1.1 | pure | 1940–1960 | Thinning every 10 years | 11 |
| D | Stutensee | STU | 49°04'12.45"N | 8°27'00.97"E | 765 | 1.8 | mixed | 1940 | Protected forest with No interventions since 2012 | 28 |
| D | Vorsenz | VOR | 49°07'16.0"N | 8°27'37.5"E | 115 | 2.7 | mixed | 1940 | Protected forest with no interventions since 2003 | 28 |
| D | Wupperthal | WUP | 51°12'00.0"N | 7°06'36.0"E | 187 | 0.5 | pure | 1980 | Initial tending and first thinning | 13 |

In order to identify the geographic origin of the introduced Oriental beech, we sampled 12 to 30 adult trees from each stand (Table 2), including both putative European and putative Oriental beech trees based on leaf morphological traits, following Denk (1999a) and Fitschen (2017). Although Oriental beech trees tend to have larger leaves (Oriental beech: 8– 17 cm, European beech: 5–10 cm), more veins (Oriental beech: 8–12, European beech: 5–8), a more elongated leaf shape and more rounded leaf base, the two subspecies cannot be distinguished with confidence based on morphological traits alone, especially in the presence of hybrid individuals (see Results).

We studied hybridization in offspring at two sites, ALL and WAE. We chose these two sites because they were mixed plantations and were among the oldest planted Oriental beech stands we could identify, being approximately 100 years old (Table 2). At WAE, six mature Oriental beech trees were located in a patch of 0.1 ha, surrounded by European beech forest to the south and by agricultural land on all other sides. The study area was approximately 3.3 ha. At ALL, around 60 adult Oriental beech trees were located in a 0.5 ha forest patch surrounded by European beech on three sides, while one side was bordered by a spruce plantation. The study area was approximately 4.5 ha.

A further factor in the choice of the two sites was that natural regeneration was abundant. Since fecundity typically increases with increasing stem dimensions (Oddou-Muratorio et al., 2018) and as natural regeneration was primarily found in clearings in the vicinity of large trees, a random or transect sampling of the offspring seemed inefficient. Instead, we opted for an adaptive sampling scheme that increased the sampling density near the largest trees, in terms of their diameter at breast height (DBH). These trees are generally also the tallest and have the largest crowns, and thus have less competition and capture more pollen, potentially not only from direct neighbors but also from more distant trees. Based on a pilot genotyping study (Kurz, 2018; Kurz et al., 2019), we chose the 3 largest Oriental and the 3 largest European beech trees and sampled offspring along 32-m cross-transects with the focal trees (i.e. the putative mother trees) at the cross point (Fig. S1). We then set up circular plots with a radius of 2 m on each transect at a distance of 2, 6 and 14 m from the focal tree (Fig. S1). In each circular plot, we counted all seedlings and saplings with a height ≤3 m, and we sampled and genotyped a fraction of them: 20% for WAE and 10% for ALL, with the exception of one tree with a high offspring density, where only 3% of them were sampled and genotyped).

### Climate data

We used downscaled historical data to characterize the climate at the sites of the sampled natural Oriental beech populations and the 11 planted stands. We used monthly minimum and maximum temperature and precipitation time series of the CHELSAcruts (1901–2016) data set (Karger et al., 2017; Karger & Zimmermann 2018). We restricted the data set to the period from 1 January 1901 to 31 December 1978. This is because starting in approximately 1980, the temperature values have been affected by global warming (Harris et al., 2014), and we wanted to assess the climatic conditions to which the natural European and Oriental beech populations have adapted prior to this change. We calculated quarterly mean temperature and summed precipitation for each year and then averaged these values across the years to obtain a single quarterly temperature and precipitation value for each sampling site. For simplicity, hereafter we refer to the first quarter (i.e. January, February, March) as winter, the second as spring, the third as summer and the fourth as fall.

### DNA isolation and microsatellite genotyping

We isolated DNA from bud, leaf or cambium tissue (Table S1), using the NucleoSpin 96 Plant Kit (Macherey-Nagel, Düren, Germany; for buds and leaves) and the E.Z.N.A. SP Plant DNA Kit (Omega Bio-Tek, Norcross, GA, USA; for cambium) and following the manufacturers’ recommendations. We used buds or leaves, depending on the season of the sampling, and cambium for the trees we could not reach with a pruner (Table S1). We used 16 microsatellite loci developed for European beech, but which are also polymorphic in Oriental beech, with 33 alleles that do not occur in European beech in the study of Lefèvre et al. (2012). We carried out the PCR amplification in two multiplex reactions. The 10 μl PCR mixture consisted of 5 μl KAPA2G Robust HotStart ReadyMix (Roche), 1 μl molecular grade water, 1.5 μl primer premix (each primer had a concentration of 1 μM), and 2.5 μl of tenfold-diluted DNA. We applied the same PCR conditions to both multiplex reactions. Initial denaturation at 95°C for 5 min was followed by 35 cycles consisting of denaturation at 95°C for 30 sec, primer annealing at 50°C for 30 sec, and extension at 72°C for 30 sec. Following the cycling, a final extension of 30 min at 72°C was added. For the subsequent genotyping on an ABI-3500 Genetic Analyzer (Applied Biosystems, USA), 0.5 μl of fivefold-diluted PCR product was mixed with 9.25 μl of Hi-Di formamide and 0.25 μl LIZ600 size standard (ThermoFisher Scientific, Waltham, MA, USA), denatured for 10 min at 95°C and then immediately cooled on ice. We used the software TANDEM (Matschiner & Salzburger, 2009) to sort alleles by raw size and to detect discrete size variants. Based on these results, we defined bins which were used in the GeneMapper software (ThermoFisher Scientific) to assign each allele to a bin.

We successfully genotyped a total of 115 individuals from natural Oriental beech populations and 213 adult trees from across the 11 planted stands. Additionally, we genotyped 245 seedlings and saplings from sites ALL and WAE to estimate the extent of hybridization between the two subspecies.

### Phenological observations

We studied the spring phenology of Oriental and European beech at WAE and LEI across two years. For these observations, we selected only previously genotyped trees, for which the pure subspecies or F1 hybrid identity had been confirmed: 6 and 10 adult Oriental beech trees in WAE and LEI and 13 and 10 adult European beech trees in WAE and in LEI, respectively. Further, in WAE, in 2021, we monitored bud development of every selected tree from 14 April to 12 May 2021 at an interval of five to ten days. In contrast, in LEI, and in WAE in 2022, we only observed the phenological status only at day of the year 115. We considered the bulk of the foliage and recorded the phenological stage reached by at least 75% of all buds using a categorical scale (Vitasse et al., 2013) comprising the following five stages: (0) dormant bud, (1) bud swelling, (2) bud burst, (3) leaf-out, (4) leaf unfolded (Fig. S2). The same persons performed the observations across years, but different persons across sites. We observed adult trees using binoculars (magnifying power: 10 × 30 with Image Stabilization technology) at a distance of approximately 15–20 m from the target tree.

### Statistical analyses

To assess patterns of genetic diversity of the sampled natural Oriental and European beech populations, we used the software Arlequin version 3.5.2.2. (Excoffier & Lischer, 2010) to estimate the number of alleles (Na), the observed and expected heterozygosity (H_O_ and H_E_), the allelic richness (Ar), the Garza-Williamson Index (M-ratio), a measure to detect population bottleneck events, and the inbreeding coefficient (F_IS_). Since the sample sizes at each individual location were small (3 to 14 individuals, see Table 1), we calculated these statistics for several sites grouped together based on mountain ranges (Table 1). While most of these groupings were obvious, the genetic clustering of the south-western Black Sea coast sites to a population became clear only after the analysis of population structure (see below). Finally, we searched for alleles that are specific to one part of the distribution range of Oriental beech, thereby refining the study of Lefèvre et al. (2012), who described new Oriental beech alleles for the subspecies as a whole.

To identify the main genetic clusters across the natural range of European and Oriental beech (Fig. 1) using data from all natural Oriental beech populations (N=115), as well as four European beech trees from each of the six admixed stands in Western Europe (N=24), we used the Bayesian clustering method implemented in the software Structure version 2.3.4 (Pritchard et al., 2000). We started from the model with correlated allele frequencies and admixture and considering models from one to ten clusters (K). We eliminated the first 100,000,000 iterations as a burn-in period, and we used the following 500,000 iterations for estimation. We ran twenty independent chains for each K and averaged them using CLUMPP (Jakobsson & Rosenberg, 2007). Subsequently, we applied STRUCTURE HARVESTER version 0.6.94 (Earl & vonHoldt, 2012) to process the different runs and to estimate the number of clusters that best explained our data, considering the log likelihood of each K (L(K) method; Pritchard et al., 2000) and the ad hoc statistic ΔΚ (Evanno method; Evanno et al., 2005). We used an analysis of molecular variance (AMOVA), implemented in Arlequin, to describe the partitioning of genetic variation into variation between clusters, within clusters and within individuals. To estimate the degree of genetic divergence between the identified genetic clusters, we calculated the pairwise F_ST_ between them using Arlequin.

To track the origin of the 11 Oriental beech stands planted in Western Europe, we used two approaches. First, we considered the natural populations used in the above model (N=139) as learning samples and used the USEPOPINFO model of Structure with K=6. This model has been successfully used in several previous studies to identify the origin of new samples (e.g., Randi et al., 2001). We assessed the sensitivity of this model to the MIGRPRIOR (0 and 0.2), to POPALPHAS (0 or 1), and to ALPHAMAX (10 and 1), where the first value indicates the default. Second, we used the genetic clustering algorithm of the Structure software in an identical way as for the natural populations (i.e., without USEPOPINFO), but simply adding the genetic data from the stands of unknown origin to the data set. Here again, we only used K=6 and 20 replicates. Since genetic clustering algorithms are known to be affected by unequal sampling (Puechmaille, 2016), we performed this analysis for each stand separately. We followed this procedure for two reasons in particular: (i) we wanted to avoid over-representation of this cluster, as preliminary analyses showed that at least WAE and ALL have a Caucasian origin, and (ii) we suspected the presence of several European beech individuals among the sampled trees from each mixed stand. For both approaches, we used the model with correlated allele frequencies, assumed six clusters, ran 20 independent chains, eliminated the first 100,000,000 as a burn-in, and used 500,000 iterations for estimation.

We tested if the two subspecies hybridize and estimated the rate of hybridization at WAE and ALL using the USEPOPINFO model implemented in Structure. We performed the analysis in two steps. We first applied the clustering algorithm with K=2 to the adult trees, which confirmed the presence of two clear clusters (see Results). Then, we applied the USEPOPINFO model, again with K=2, to the offspring to estimate the proportion of their Oriental beech ancestry. We ran separate analyses for WAE and ALL. Other settings and post-processing of the runs were identical to the previously described analyses. We used ancestry coefficients (q) from the structure software to classify individuals to the closest plausible level of hybridization. The WAE stand was established in 1921 and the ALL stand in 1923; with a generation time of 30 to 40 years (Kandemir & Kaya 2009), we thus expected that hybridization could be ongoing for two to three overlapping generations. If a cross between two mature F1s, which would lead to F2s, has a small probability, the most common levels of hybridization should be: purebred, F1 and backcross. Thus, we decided to bin the ancestry coefficients (q) from Structure into the closest category among purebred (Oriental beech if q=1 and European beech if q=0), F1 (q=0.5), and backcross (q=0.75 or 0.25), where q is the proportion of Oriental beech ancestry.

Finally, to assess phenological differences between the two subspecies, in WAE, in 2021, we estimated the day of year for each stage for the two subspecies using a linear interpolation between the observed stages on two consecutive observation dates. For LEI and WAE in 2022, we compared the phenological stages using a t-test.

## Results

### Climate at the natural sites and planting sites

Oriental beech growing sites varied widely in their climate, reflecting the wide geographic range of the sampled populations (Fig. 1a and c, Table S2). The site in the Elburz Mountains in Iran was warmest (summer and winter mean temperature as high as 25°C and 7°C, respectively), while growing sites in the Greater Caucasus and the easternmost population of the Pontic Mountains were coldest, with mean winter temperature as low as - 6°C in GOD and at most 1°C at the warmest sites. Growing sites in the Greater Caucasus had variable precipitation regimes, with as little as 608 mm of total annual precipitation in ING (Table S2). The site from the Taurus Mountains (HAT) had a very continental climate: mean temperature varied from 3°C in winter to 21°C in the summer, while the total annual precipitation was as high as 1085 mm (Fig. 1c, Table S2). The Pontic mountain growing sites became warmer and drier from east to west. They were among the most humid sites, with total annual precipitation as high as 1200 mm (ORD) and at least 1015 mm (KAS) (Table S2). The two sites from the south-western Black Sea coast in Bulgaria (KOS and NES) were the driest and warmest sites, with only 519 and 670 mm of total annual precipitation (in comparison to 838 mm in NOO, Table S2).

The sites in Western Europe were warmer than the coldest Oriental beech sites (Greater Caucasus) and were characterized by a low degree of seasonality, especially in terms of precipitation, in comparison to the Oriental beech growing sites (Fig. 1c). We estimated precipitation seasonality as the standard deviation of the monthly precipitation estimates divided by the annual mean precipitation, averaged across the years, which was, on average, 0.58 for natural Oriental beech sites (range: 0.41–0.78) and 0.46 for the sites in Western Europe (range: 0.44–0.54). The coldest and wettest site of the latter group was the Swiss site LEN, while the sites in the Rhine valley were the hottest and driest (ALL, LEI, STU, VOR, RIE; Fig. 1 and Table S2).

### Genetic diversity in European and Oriental beech

All individual Oriental beech clusters had a higher genetic diversity than European beech in terms of expected heterozygosity (H_E_), except the Elburz Mountains cluster (Table 3; note that mountain range/region corresponds to the genetic cluster). All populations had a slight heterozygote deficit, i.e., the observed heterozygosity (H_O_) was lower than the expected heterozygosity (H_E_). Depending on the cluster, up to four loci (25%) deviated significantly from the Hardy-Weinberg equilibrium, which was due to the presence of four extremely polymorphic loci in the data set (FS1_15, DZ447, sfc_1143, EMILY; Table S3 and Fig. S3). Furthermore, all Oriental beech clusters together had, on average, 14.4 alleles per locus, compared with 6.6 alleles per locus in European beech, even though the individual Oriental beech clusters had at most only 10.3 alleles (south-western Black Sea coast) and as few as 6.2 alleles (Elburz Mountains; Tables 3 and S3, Fig. S3). The mean allele size range was also greater in Oriental beech (29.5) than in European beech (17.25). The allele size range of Oriental beech clusters was consistently large. Thus, even though Oriental beech had more alleles than European beech, they still had fewer than expected based on their allele size range, suggesting population size reduction in the past (see M-ratios in Table 3).

**Table 3.** Summary of genetic diversity in natural European and Oriental beech populations estimated from 16 nuclear microsatellite loci. Summaries are provided for the six genetic clusters, based on the mountain ranges and refined by the genetic clustering (see Figure 1), averaged across all loci. N: number of individuals genotyped; k: mean number of alleles; r: allele size range; Ar: allelic richness; M-ratio: Garza-Williamson Index calculated as M = k/r, where smaller values indicate a bottleneck event; H_O_: observed heterozygosity; H_E_: expected heterozygosity; HWE%: percentage of loci that significantly deviated from the Hardy-Weinberg equilibrium; F_IS:_ inbreeding coefficient with p-values based on a permutation test, with ns indicating non-significant.

| Mountain range/<br>Region | Subspecies | N | k | r | Ar | M-ratio | $H_o$ | $H_E$ | HWE% | $F_{IS}$ |
| --- | --- | --- | --- | --- | --- | --- | --- | --- | --- | --- |
| Elburz Mountains | Oriental beech | 14 | 6.2 | 17.6 | 5.69 | 0.35 | 0.67 | 0.67 | 6 | -0.0235 (ns) |
| Greater Caucasus | Oriental beech | 34 | 9.3 | 23.9 | 6.3 | 0.39 | 0.69 | 0.73 | 25 | 0.0588 (0.01) |
| Taurus Mountains | Oriental beech | 14 | 6.4 | 19.3 | 6.1 | 0.33 | 0.7 | 0.76 | 12 | 0.0350 (ns) |
| Pontic Mountains | Oriental beech | 25 | 9.4 | 25.9 | 6.96 | 0.36 | 0.69 | 0.77 | 25 | 0.0725 (0.008) |
| SW Black Sea coast | Oriental beech | 28 | 10.3 | 24.2 | 6.93 | 0.43 | 0.65 | 0.77 | 25 | 0.1126 (<0.001) |
| Western Europe | European beech | 24 | 6.6 | 15.1 | 5.47 | 0.44 | 0.67 | 0.71 | 19 | 0.0356 (ns) |

Lefèvre et al. (2012) already demonstrated the transferability of the 16 microsatellite loci used herein for Oriental beech, but only for a limited number of samples of unknown origin. We found that all loci were polymorphic across all clusters (Tables 3 and S3), demonstrating the transferability of these loci across the whole range of Oriental beech. Additionally, we identified several private alleles for each cluster, with the south-western Black Sea coast cluster having the most (15), the Elburz Mountains and the Greater Caucasus clusters having 9 and 8, respectively, and the Taurus Mountains having just 4 (Table S4).

### Genetic clusters in European and Oriental beech

The hierarchical AMOVA indicated that most of the variance in allele frequencies was within clusters and individuals, even though a strong and significant genetic structure was detected among mountain ranges/regions (F_ST_=0.1378, p<0.001; Table S5). The most genetically differentiated parts of the distribution range were also the most geographically distant ones, i.e., between European beech from Western Europe and Oriental beech from the Elburz Mountains (pairwise F_ST_=0.25) or the Greater Caucasus (pairwise F_ST_=0.21). The divergence between the two subspecies dropped to 0.15 (pairwise F_ST_) between European beech and Oriental beech from the south-western Black Sea coast (Table 4).

**Table 4.**
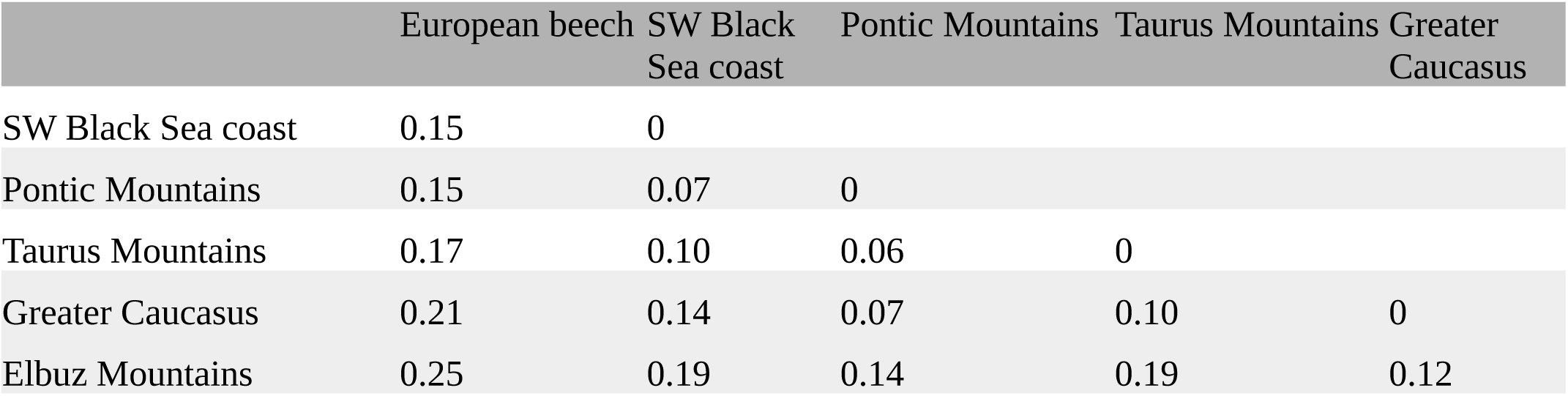
Pairwise F_ST_ among the six genetic clusters. All values are significant at the 0.05 level.

Genetic clustering assuming different numbers of clusters (K) led to a separation of European and Oriental beech at K=3, while at K=2 Oriental beech from the south-western Black Sea coast clustered with European beech (Fig. S4). Higher K values improved the model fit and resulted in additional clusters within the range of Oriental beech, while all six European beech sites clustered together (Fig. S4). We found that a model assuming K=6 provided a good fit to the data using various criteria, including L(K) and Evanno’s method (Figs. S4 and S5), with only a few individuals having admixed origin (Fig. 1b). Further, this clustering corresponded to the distinct mountain ranges, except for the south-western Black Sea coast region, which is not a mountain range but rather a set of loosely connected populations situated in Turkey and Bulgaria (Table 1, Fig. 1a and b). It appears that this latter cluster is separated from the neighboring Pontic Mountain cluster, and their contact zone is somewhere between the Turkish towns of Inebolu (KAS) and Zonguldak (ZON, Fig. 1a and b).

### The origin of planted Oriental beech in Western Europe

The assignment of the origin using the two methods, i.e., with and without USEPOPINFO, led to similar conclusions, with slightly fewer clear assignments when the USEPOPINFO model was used (Figs. 2 and S6). This is probably because (i) the assignment with USEPOPINFO is more difficult with more than two clusters (no literature exists for the performance in such cases), (ii) the six clusters are not entirely pure (presence of some admixed individuals), and (iii) some of the clusters are not in Hardy-Weinberg equilibrium at all loci. Using USEPOPINFO replicate chains led to almost exactly the same assignment of individuals. MIGRPRIOR and ALPHAMAX did not influence the assignment, while POPALPHAS=0 resulted in a less clear assignment, namely, with all individuals sharing some ancestry from all clusters (not shown).

**Figure 2.**
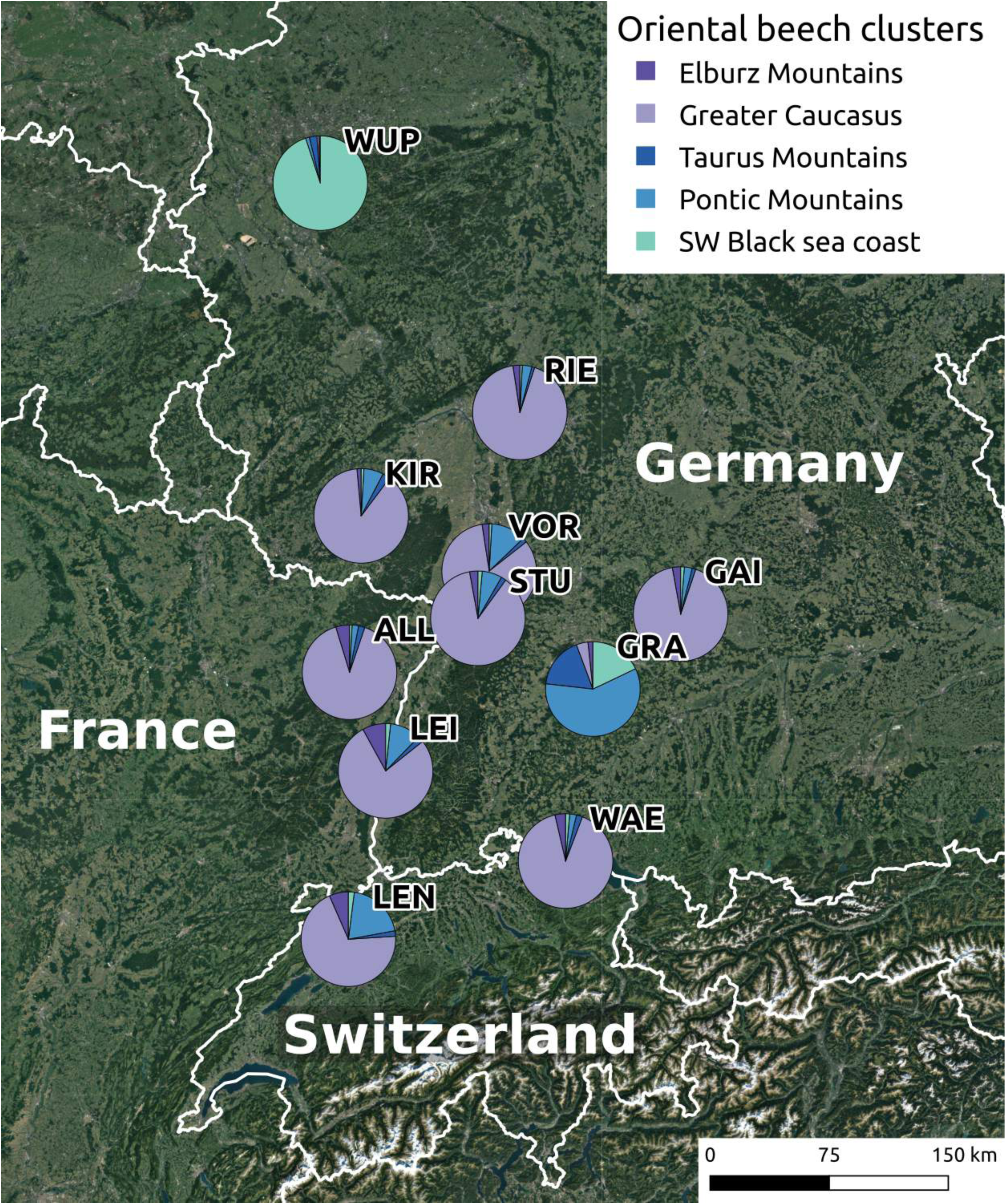
Origin of Oriental beech in 11 Western European plantations. Each pie chart shows the proportion of the five Oriental beech clusters in the gene pool of all sampled Oriental beech trees.

Several adult trees from each stand were sampled based on morphological traits and without knowledge of their true subspecies identity. Genotyping indicated that some trees had been assigned to the wrong subspecies based on morphological traits, as well as potential hybrids and backcrosses among these adult trees. The planted Oriental beech trees from nine out of eleven stands originated from the Greater Caucasus (Fig. 2), including the two Swiss sites, the one French site, and six of the German sites (GAI, KIR, LEI, RIE, STU, VOR). Most individuals had a small proportion of ancestry from other Oriental beech clusters, most often from the neighboring clusters such as from the Pontic and Elburz Mountains. LEN had the highest proportion of foreign ancestry, suggesting a possible origin from the southern slopes of the Greater Caucasus or the easternmost part of the Pontic Mountains. The plantation WUP was clearly assigned to the south-western Black Sea cluster, while the plantation GRA could not be clearly assigned to any of the clusters but had the highest proportion of ancestry from the Pontic Mountains (Fig. 2).

### Hybridization at two planting sites

Genetic clustering indicated that the probability of the data was highest at K=2 both at WAE and at ALL (results not shown). Across the two sites, ancestry coefficients from the Structure software suggested that European and Oriental beech hybridize extensively. The proportion of hybrid or backcross seedlings and saplings were, on average, 41.3% in WAE and 17.8% in ALL, but with a large variation in space (between 0 and 91% in WAE and 0 to 36% in ALL). However, the proportion of putative F1 and backcrosses and their spatial distribution differed between the two sites. At WAE, 68 (64.8%) offspring were classified as putative purebred European beech and three (2.8%) as Oriental beech, while 21 (20%) and 13 offspring (12.4%) were classified as putative F1 and backcross, respectively (Table 5, Fig. S7). All putative F1 offspring were found beneath the six adult Oriental beech trees (Fig. 3). Two seedlings found under crowns of adult European beech trees were classified as putative backcross to a pure European beech tree (i.e. Oriental beech ancestry close to 0.25), while 11 other offspring found under crowns of adult Oriental beech trees were backcrosses to Oriental beech (Figs. 3 and S7). At ALL, 95 (67.9%) of all genotyped seedlings and saplings were classified as pure European beech and 25 (17.9%) as pure Oriental beech. Eleven individuals (7.9%) were putative F1s (Table 5, Fig. S7), but, in contrast to WAE, they were distributed throughout the whole study area, in the vicinity of both adult European and Oriental beech (Fig. 3). Nine offspring (6.4%) could be classified as putative backcross to a pure European beech tree and two (1.4%) to a pure Oriental beech tree (Table 5, Fig. S7).

**Figure 3.**
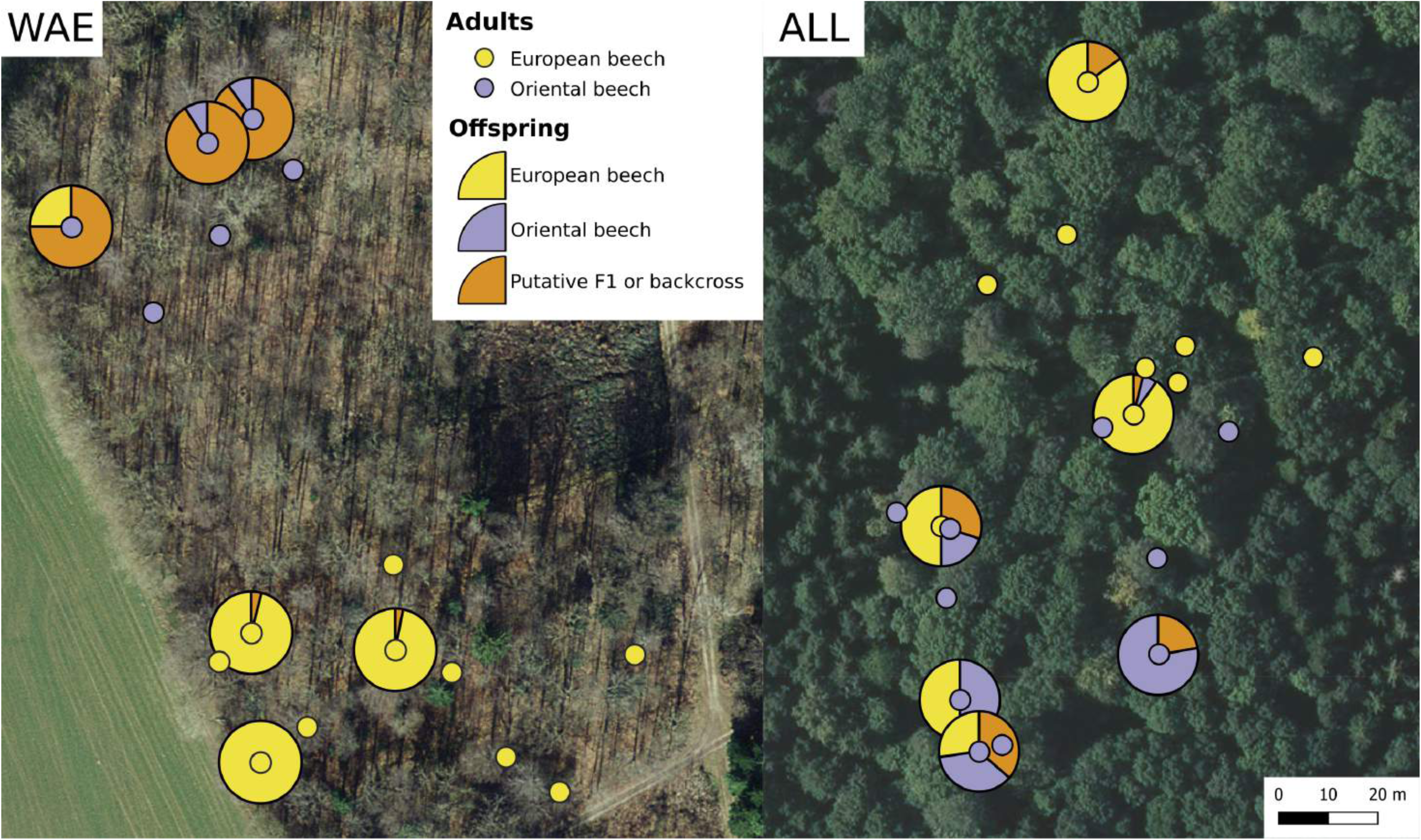
Evidence for extensive hybridization between introduced Oriental beech and native European beech in two plantations **(a)** Waldi (WAE) and **(b)** Allenwiller (ALL). Circles show the sampled adult trees, while pie charts show the proportion of European and Oriental beech ancestry for seedlings around them. A total of 105 seedlings from Wäldi and 140 from Allenwiller were genotyped.

**Table 5.** Hybridization at two sites: Waldi (WAE) and Allenwiller (ALL). Seedlings and saplings up to 3 m in height were counted around the six largest adult trees. Their total number is indicated by N (t), of which N (g) were genotyped. See Materials and Methods for more details and Fig. S1 for the sampling configuration. The putative status is derived from the estimated ancestry coefficients using the software Structure version 2.3.4 (Pritchard et al., 2000) with K=2 by binning them to the closest biologically plausible category among purebred (Oriental beech if q=1 and European beech if q=0), F1 (q=0.5), and backcross (BC; q=0.75 or 0.25), where q is the proportion of Oriental beech ancestry. The hybridization rate is calculated for each mother tree and includes both putative F1s and backcrosses.

| Site | Mother tree |  | Number of seedlings |  | Putative status |  |  |  |  | Hybridization rate (%) |
| --- | --- | --- | --- | --- | --- | --- | --- | --- | --- | --- |
|  | Subspecies | ID | N (t) | N (g) | European beech | Oriental beech | F1 | BC to European beech | BC to Oriental beech |  |
| WAE | Oriental beech | OA001 | 103 | 20 | 0 | 2 | 13 | 0 | 5 | 90 |
| WAE | Oriental beech | OA002 | 50 | 11 | 0 | 1 | 6 | 0 | 4 | 91 |
| WAE | Oriental beech | OA003 | 22 | 5 | 1 | 0 | 1 | 0 | 2 | 60 |
| WAE | European beech | A123 | 129 | 25 | 24 | 0 | 0 | 1 | 0 | 4 |
| WAE | European beech | A144 | 170 | 33 | 32 | 0 | 0 | 1 | 0 | 3 |
| WAE | European beech | A151 | 58 | 11 | 11 | 0 | 0 | 0 | 0 | 0 |
| ALL | Oriental beech | VA015 | 16 | 4 | 2 | 2 | 0 | 0 | 0 | 0 |
| ALL | Oriental beech | VA020 | 41 | 11 | 3 | 4 | 4 | 0 | 0 | 36 |
| ALL | Oriental beech | VA044 | 58 | 18 | 0 | 14 | 3 | 0 | 1 | 22 |
| ALL | European beech | VA007 | 101 | 10 | 5 | 2 | 2 | 0 | 1 | 30 |
| ALL | European beech | VA055 | 492 | 51 | 46 | 3 | 1 | 1 | 0 | 4 |
| ALL | European beech | VAN077 | 1561 | 46 | 39 | 0 | 1 | 6 | 0 | 15 |

### Spring phenology at two planting sites

We found that adult Oriental beech trees had an earlier spring phenology than adult European beech trees across both sites and years (Fig. 4). Using the full sequence of observations from WAE in 2021, we found that the magnitude of difference between adult trees of Oriental and European beech changed across the phenological stages. Stage 2 (bud burst) was reached on average four days earlier in Oriental beech (t = 2.00, df = 21.55, p-value = 0.0579), stage 3 (leaf-out) five days earlier (t = 3.28, df = 21.64, p-value = 0.0035), and leaf unfolding (stage 4) 3.8 days earlier (t = 2.75, df = 19.54, p-value = 0.013). When focusing on comparing the average stage at day of the year 115, again at both sites and years we found that Oriental beech was earlier. In WAE, Oriental beech was, on average, in stage 2.7, while European beech was in stage 1.9 (t = 2.756, df = 20.449, p-value = 0.012) in 2021, and in stage 3.9 and in stage 2.6 (t = 4.164, df = 8.0661, p-value = 0.003) in 2022. In LEI, again on day of the year 115, Oriental beech was, on average, in stage in 3.5 and European beech in stage 1.9 (t = 4.5293, df = 16.032, p-value = 0.0003) in 2021, and in stage 4.0 and in stage 2.8 (t = 2.9599, df = 10, p-value = 0.0143) in 2022 (Fig. 4).

**Figure 4.**
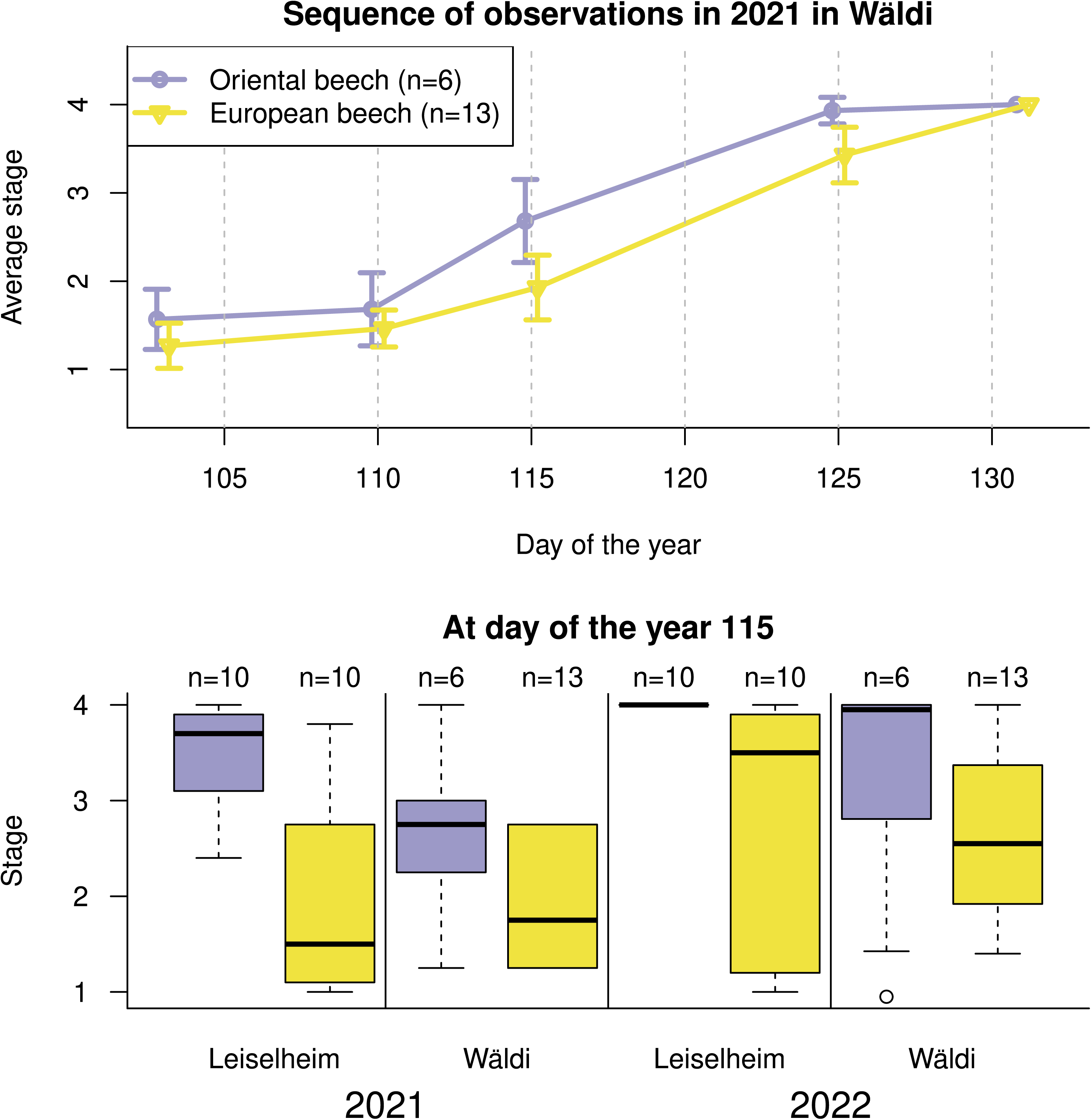
Spring phenology of introduced Oriental beech and native European beech in Waldi (WAE) in 2021 and 2022, and in Leiselheim (LEI) in 2022. Stage (0) means dormant bud, (1) bud swelling, (2) bud burst, (3) leaf-out, (4) leaf unfolded.

## Discussion

Here, we presented a database with genetic and climatic information about 11 planted Oriental beech stands in Western Europe that can serve as natural laboratories of AGF in European beech forests (Fig. 1). Using a genetic clustering algorithm and samples from across the natural range of Oriental beech, we traced the origin of the planted Oriental beech stands and found that nine of them had a Greater Caucasus origin, while two were most likely established from Turkish seed lots. Climate time series data suggested that the seed translocation involved a shift from a continental climate to conditions with less seasonal climatic variation. At the two sites, WAE and ALL, we found evidence for extensive hybridization between Caucasian Oriental beech and European beech, which might be asymmetric. The combined results of this study provide an basis for the evaluation of benefits and risks of AGF for Oriental beech to native European beech forests.

### Oriental beech for AGF: an obvious gain in genetic diversity

Consistent with previous findings based on isozymes (Gömöry et al., 2007, 2010), cpDNA sequences (Vettori et al., 2004) and microsatellites (Magri et al., 2006), we found that Oriental beech populations harbor a great amount of genetic diversity (Table 3). Genetic diversity was particularly high in the Pontic Mountains and south-western Black Sea coast clusters, where we also identified the largest numbers of private alleles (Table S4). Since our sampling of European beech was restricted to a relatively small part of the subspecies range, we cannot conclude that genetic diversity of Oriental beech is overall higher than that of European beech. Nevertheless, using the same 16 microsatellite loci, de Lafontaine et al. (2013) found genetic diversity values in European beech populations in France from both refugial and recolonized areas that were similar to those observed herein (mean Ar=6.3, H_O_=0.702 and H_E_=0.69; see our values in Table 3). In addition, we found that many alleles of Oriental beech are different from those of European beech (Fig. S3). Thus, upon introgression to a European beech forest, genetic diversity would necessarily increase. The highest gain in allelic richness would be achieved by introducing Oriental beech from the south-western Black Sea coast or the Pontic Mountains: allelic richness (Ar) of our European beech samples (5.47, Table 3) would increase to 7.19 and 7.03, respectively. The gain would be slightly smaller for the more distant, Greater Caucasus or Elburz Mountains clusters, with predicted Ar values of 7.09 and 6.86, respectively.

### Provenance matters: Which Oriental beech lineage for AGF?

Oriental beech is a genetically non-homogeneous subspecies. Therefore, the question is not whether Oriental beech is suitable for AGF, but rather which Oriental beech lineage, if any, would be most suitable. Our study illustrates that different provenances of Oriental beech have a wide range of divergence from the studied European beech populations (Table 4). Both past studies and the present investigation illustrate that Iranian Oriental beech, growing today along the Elburz Mountains, appears to be the most strongly diverged provenance from European beech, but also from the rest of the Oriental beech range. Using 12 isozyme loci and 279 populations of Oriental and European beech, Gömöry and Paule (2010) showed that Iranian beech separated from other Oriental beech populations with the deepest split within the subspecies, and it also appears to be the ancestral lineage (Gömöry et al., 2018). Bijarpasi et al. (2020) reported G_ST_ values as high as 0.503 between Iranian and European beech at expressed sequence tag-simple sequence repeats (EST-SSR), while Cardoni et al. (2021) even suggested, based on the 5S rRNA gene region, that Iranian beech is a separate species. In the present study, using nuclear microsatellites, we also found the highest pairwise F_ST_ (0.25) between Iranian Oriental beech and European beech, but a separate Iranian cluster appeared only at K=5 (Fig. S4). Even though our sampling covered the species range well, our data are not suitable alone for revising the species taxonomy, given the high mutation rate of microsatellite loci. Nevertheless, our data do show a continuous isolation-by-distance pattern from east to west (Table 4) and suggest that Greater Caucasian populations, which were not sampled by Bijarpasi et al. (2020) and Cardoni et al. (2021), provide an intermediate type between Oriental beech from Iran and the Pontic Mountains (F_ST_=0.12 and 0.14, respectively; Table 4).

The south-western Black Sea coast cluster, composed of populations sampled in north-western Turkey and eastern Bulgaria, showed the lowest divergence from European beech and they even clustered together at K=2 (Fig. S4). In Turkey, this cluster forms a contact zone with the Pontic Mountains cluster, likely situated between the populations KAS and ZON (Fig. 1a and b). The location of this contact zone appears to be consistent with findings on other tree species and historical climate data (Krebs et al., 2004). For example, Mattioni et al. (2013) suggested the presence of different glacial refugia in present-day Georgia and in the area near the Dardanelles strait in *Castanea sativa* L., which places the contact zone in the same area as in our study. The genetic proximity of the south-western Black Sea coast cluster to European beech could indicate that European beech lineages colonized the European continent from refugia spanning across the southern Balkan and western Turkey; a region well known for extremely high plant diversity and source for westward range expansions (e.g. Bagnoli et al., 2016; Feliner, 2014; Mansion et al., 2009). Indeed, Gömöry et al. (1999, 2007) suggested that the Balkan peninsula contains a mixture of European and Oriental beech gene pools. More recently, Cardoni et al. (2021) suggested, based on the 5S rRNA gene region, that Oriental beech from Greece occurs in a recent contact zone, and that European beech has a potential hybrid origin. Such an evolutionary history would be no exception. Indeed, up to 25% of plant species are thought to be of hybrid origin, and genetic admixture of divergent lineages is known to play an important role in the success of colonizing populations (Mallet, 2005; Rius & Darling, 2014; Soltis & Soltis, 2009). Although we did not sample in Greece, our data from the Balkans revealed that F_ST_ was relatively high between the south-western Black Sea coast and European beech clusters (0.15), while the south-western Black Sea coast populations had an F_ST_ of 0.07 and 0.10 with the Pontic Mountains and Taurus Mountains clusters, respectively (Table 4), suggesting that our populations from eastern Bulgaria are closer to Turkish Oriental beech. Nevertheless, we note that F_ST_ between European beech and south-western Black Sea coast populations could have also been inflated by the low diversity in European beech (Charlesworth, 1998).

The strong genetic clustering within the range of Oriental beech indicates that each of the clusters should be assessed experimentally before AGF can be recommended, especially because the risk of outbreeding depression increases with genetic distance between host and donor populations (Aitken & Whitlock, 2013; Frankham et al., 2011). There are three primary mechanisms of outbreeding depression: chromosomal incompatibility, breakdown of co-adapted gene complexes (epistatic interactions), and local adaptation losses resulting from the introduction of locally maladapted alleles (Aitken & Whitlock, 2013; Edmands, 2006; Whitlock et al., 2013). Ribeiro et al. (2011) found two paralogous 5S rRNA loci in *Fagus sylvatica*, suggesting chromosomal rearrangements during its evolution. Cardoni et al. (2021) also found two 5S rRNA clusters, and these separate the phylogenetically distinct A and B lineages in the *F. sylvatica* species complex. The B-lineage successively replaces the A-lineage along a south-east (Oriental beech, Iran) to north-west (European beech, Germany) gradient. Thus, there might be some degree of chromosomal incompatibility between Iranian Oriental beech and European beech, but further genomic/experimental evidence is needed. The other mechanisms are even more difficult to pin down. This is because outbreeding depression rarely occurs already in the F1 generation (Goto et al., 2011), as it takes generations for recombination to break up beneficial gene interactions. For example, line-cross analysis of multiple hybrid (F1, F2 and backcrosses) and pure-species populations of two diploid eucalypt species, *Eucalyptus globulus* and *Eucalyptus nitens*, showed that F2s and backcrosses showed stronger outbreeding depression than F1s in survival and growth, and that this effect increased with increasing age (Costa e Silva et al., 2012). Based on pairwise F_ST_ between distinct Oriental beech clusters and European beech, we can only suspect that the risk of outbreeding depression is highest when Iranian Oriental beech introgresses to European beech.

### Hybridization rate: detectability and spatial configuration

Nine of the eleven planted Oriental beech stands had a Greater Caucasian origin, including the two stands at WAE and ALL, where we also detected extensive hybridization among the offspring growing in proximity to dominant beech trees (Fig. 3, Table 5). According to the simulation study of Vähä and Primmer (2005), the divergence between Caucasian Oriental beech and European beech (F_ST_=0.21) and 16 polymorphic microsatellite loci were sufficient for distinguishing between purebreds and F1s with nearly 100% accuracy. Identification of backcrosses was less accurate (80%). With lower divergence between host and donor populations, the detection accuracy would drop given the same 16 microsatellite loci used herein. Thus, future AGF programs with provenances that are genetically closer to European beech would require a larger number of loci for tracking hybridization.

With an average hybridization rate of 31.4 (WAE) and 17.8% (ALL), including both putative F1s and backcrosses, we conclude that the hybridization rate is high (Fig. 3, Table 5) and is similar to that found in other species that hybridize extensively, such as European white oaks (Gerber et al., 2014), poplars (Talbot et al., 2012), and larches (Meirmans, 2019). Forest management might also explain the high rates of hybridization. Both sites have even-aged management, and thinning can favor the growth of new seedlings and thus the growth and propagation of hybrid individuals. Indeed, it has been widely recognized that human-impacted landscapes can facilitate hybridization (Hoban et al., 2016), including the propagation of invasive species (Radtke et al., 2013), but also as a management tool to facilitate it (Grabenstein & Taylor, 2018). Unfortunately, however, we do not have information about the management schemes for these sites during the past 100 years.

The spatial distribution of hybrids showed a sharp contrast between the two sites (Fig. 3). At site WAE, below the three studied Oriental beech trees we found almost exclusively F1 offspring, while below the three studied European beech trees we found almost exclusively pure European beech offspring (Fig. 3, Table 5). Indeed, the six Oriental beech trees at this site are surrounded on two sides by numerous European beech trees, and by an agricultural field on the other sides. Thus, we may speculate that the local pollen cloud is dominated by European beech, leading to asymmetric hybridization in the direction of Oriental beech and explaining the near complete lack of pure Oriental beech seedlings (Fig. 3). Evidence for asymmetric pollen flow from European beech to Oriental beech is also supported by the fact that putative backcrosses were only found the direction of Oriental beech based on ancestry coefficients (Table 5). In contrast, at site ALL, the rate of hybridization was more equally distributed among Oriental and European beech mother trees, ranging from low (0% or 4%) to intermediate (30% or 36%) in both subspecies (Fig. 3, Table 5). Although the spatial configuration is similar to WAE, i.e. the group of adult Oriental beech trees is surrounded on two sides by numerous European beech trees, there are also at least 55 adult Oriental beech trees, which increase the frequency of conspecific pollen. Such patterns of frequency-dependent asymmetric hybridization have been observed in oaks (Lagache et al., 2013). Further, using simulations, Klein et al. (2017) showed that grouped individuals can protect themselves from hybridization, more so than randomly distributed individuals. This would also explain the high proportion of hybrids in the center of our study area at ALL, where the two species are mixed (Fig. 3). Such information could help improve the design of effective AGF programs.

### Climatic and phenological decoupling

Assisted gene flow in forest tree species is often justified by the climatic decoupling between locally adapted tree populations and the past environmental conditions under which they have evolved (Sáenz-Romero et al., 2020). Indeed, Frankham et al. (2011) emphasized that the ability of a species to occupy substantially different environments can be useful for predicting the risk of outbreeding depression. Our climate data from the growing sites of the studied natural populations and the planted stands in Western Europe suggest that the climatic envelope of all Oriental beech populations is wide (Fig. 1c). When Oriental beech from the Caucasus is moved to the Western European sites, it is shifted to a climate with less seasonal variation in temperature and precipitation (Fig. 1c, Table S2). The annual average temperature and precipitation of other Oriental beech clusters that are genetically closer to European beech were more similar those of the planted sites (Table S2), but their precipitation seasonality was different (Fig. 1c).

The effectiveness of AGF will depend not only on the climatic match but also, as a consequence, on the phenological match between host and donor populations (Wadgymar & Weis, 2017). Differences in flowering phenology are considered an important reproductive barrier between closely related species (Rieseberg, 1997, 2001). At WAE, we observed that most hybrid individuals were detected in the vicinity of Oriental beech trees, which may suggest asymmetric hybridization (Fig. 3, Table 5). The potential causes for this finding are numerous and involve both pre- and postzygotic reproductive barriers (Widmer, Lexer, & Cozzolino, 2009). Our observations of spring leaf phenology show that bud break of Oriental beech precedes that of European beech and this result is consistent across years and sites (Fig. 4). Using the sequence of observations from WAE in 2021, we estimated that Oriental beech is, on average, four days earlier, however, this finding appears conservative given that the difference in LEI, and in 2022 at day of the year 115 was even larger (Fig. 4). Our observations are also consistent with previous observations in a planted Oriental beech stand in Germany (Moosmayer, 1958). Since the timing of flowering is tightly coupled with leaf unfolding in beech (Nielsen & Schaffalitzky de Muckadell, 1953), flowering asynchrony between European and Oriental beech could explain the asymmetric hybridization. In beech, full pollen dispersal typically occurs a few days after female flowers reach receptivity (Nielsen & Schaffalitzky de Muckadell, 1953). This sequence of flowering events, together with earlier flowering in Oriental than in European beech, however, indicates hybridization in the direction of European beech and not vice versa. At WAE, only six adult Oriental beech trees grow today, surrounded by many European beech trees. Therefore, it seems more likely that pollen limitation causes asymmetric hybridization. This is supported by the fact that very few purebred Oriental beech seedlings, but many hybrid seedlings, were detected, despite the six adult Oriental beech trees growing in a small pure group (Fig. 3). At ALL, where hybrid offspring were detected across the entire study site, the density of adult Oriental beech trees was larger (Fig. 3). Patterns of abundance-dependent asymmetric hybridization have also been observed in oaks (Lepais et al., 2009), and they highlight the importance of the number and spatial distribution of introduced trees when designing effective AGF programs.

### Future research needs

European beech is strongly affected by climate change: changes to its growth, reproduction, regeneration and mortality have all been linked with the warmer and drier climate, particularly with extreme hot-drought events (Hacket-Pain et al., 2016; Nussbaumer et al., 2020; Peñuelas et al., 2007), resulting in a substantial reduction of its productivity (Reyer et al., 2014; Trotsiuk et al., 2020). The potential advantage of introducing southern European beech provenances has been assessed by many previous studies, however, there are no substantial differences in genetic diversity across the range (e.g., de Lafontaine et al., 2013) and limited evidence for local adaptation and strong evidence for plasticity (e.g., Gárate-Escamilla et al., 2019; Kurjak et al., 2019; Muffler et al., 2021; Sáenz-Romero et al., 2019), suggesting that the benefits of such provenance movements appear limited. In contrast, the introduction of Oriental beech has a high potential to increase genetic diversity, and more drought-resistant provenances could also introgress pre-adapted alleles to European beech populations, thus promoting the evolutionary response of European beech to climate change. The degree of maladaptation resulting from climate change may not be homogeneous across the species range. For example, several studies have indicated strong growth reduction at the southern range edge of European beech (Jump et al., 2006; Peñuelas & Boada, 2003; Piovesan et al., 2007; Schuldt et al., 2020) but also at lower elevations in the center of its range (Dulamsuren et al., 2017; Härdtle et al., 2013). However, the extent to which different growth-determining factors, such as temperature, water and N availability and their interaction, affect European beech under different site conditions remains unclear. In this context, a network of existing Oriental beech stands growing under a wide range of site conditions provides excellent case studies for evaluating the drought and heat tolerance of the two subspecies growing side by side, along with their hybridized offspring. For this purpose, dendroecological and wood anatomical studies (of adult trees) in combination with ecophysiological methods (for studying offspring) appear to be promising approaches to provide the scientific basis for the potential introduction of Oriental beech into European beech forests. We invite other researchers across Europe to contribute with any known additional sites with existing Oriental and European beech stands to this network.

## Data Availability Statement

The data that support the findings of this study will be made openly available in a public depository upon acceptance.

## Acknowledgments

This work was supported by the following funding agencies and grants: financial support from the Federal Office for the Environment (FOEN) awarded to CS, KC and PB, a WSL Internal Innovative Project grant awarded to CS and KB, financial support from the Department of Life Sciences and Facility Management of the Zurich University of Applied Sciences (ZHAW) in Wädenswil, Switzerland) awarded to FR and THMS, the ERC Consolidator Grant “MyGardenOfTrees” awarded to KC, and financial support from the Otto Henneberg-Poppenbüttel Foundation awarded to the Chair of Silviculture at Freiburg University. We are grateful to Dr. Bertram Leder, Ralf-Volker Nagel, and Bernhard Mettendorf for locating and facilitating access to the Oriental beech stands in Germany. We thank Samuel Gunz and Roger Köchli for their contributions to the field work and Marco Walser, Rene Graf and Nicola Rhyner for help in the lab.

## List of Supplementary Material

### Supplementary Tables

Table S1. Sampling information

Table S2. Raw climate data summaries

Table S3. Number of alleles and range by loci

Table S4. Private alleles in European and Oriental beech

Table S5. Analyses of molecular variance

### Supplementary Figures

Fig. S1. Sampling scheme

Fig. S2. Definitions of *Fagus* phenological stages

Fig. S3. Allele frequency distributions per cluster

Fig. S4. Genetic clustering analyses with K from 2 to 6

Fig. S5. Summary statistics for the performance of the genetic clustering

Fig. S6. Origin of Oriental beech in 11 Western European plantations (depicted in Figure 2), but using an assignment test

Fig. S7. Ancestry coefficients for genotyped offspring in the sites WAE and ALL

