## Supplementary Tables and Figures for "Tracing the origin of Oriental beech stands across Western Europe and reporting hybridization with European beech – implications for assisted gene flow"

### List of Supplementary Material

#### Supplementary Tables

Table S1. Sampling information

Table S2. Raw climate data summaries

Table S3. Number of alleles and range by loci

Table S4. Private alleles in European and Oriental beech

Table S5. Analyses of molecular variance

#### Supplementary Figures

Fig. S1. Sampling scheme

Fig. S2. Definitions of *Fagus* phenological stages

Fig. S3. Allele frequency distributions per cluster

Fig. S4. Genetic clustering analyses with K from 2 to 6

Fig. S5. Summary statistics for the performance of the genetic clustering

Fig. S6. Origin of Oriental beech in 11 Western European plantations (depicted in Figure 2), but using an assignment test

Fig. S7. Ancestry coefficients for genotyped offspring in the sites WAE and ALL

**Table S1.** Material collected for molecular analyses (i.e. plant tissue used for DNA extraction) and details of sampling.

| Country | ID | Material | Sampling Date | Sample collector |
|---|---|---|---|---|
| Iran | NOO | leaf | Before 2019 | Ole Kim Hansen |
| Georgia | LAG | leaf | Before 2019 | Andreas Rudolf |
| Georgia | GOM | leaf | Before 2019 | Andreas Rudolf |
| Russia | ING | leaf | Before 2019 | Ladislav Paule |
| Russia | LAN | leaf | Before 2019 | Ladislav Paule |
| Turkey | HAT | leaf | Before 2019 | Hakan Sevik |
| Georgia | GOD | leaf | Before 2019 | Andreas Rudolf |
| Turkey | ORD | leaf | Before 2019 | Hakan Sevik |
| Turkey | KAS | leaf | Before 2019 | Hakan Sevik |
| Turkey | ZON | leaf | Before 2019 | Hakan Sevik |
| Turkey | BUR | leaf | Before 2019 | Hakan Sevik |
| Bulgaria | KOS | leaf | Before 2019 | Petar Zhelev |
| Bulgaria | NES | leaf | Before 2019 | Petar Zhelev |
| Switzerland | LEN | leaf | Before 2019 | Katalin Csillery |
| Switzerland | WAE | leaf/bud | 2019-2021 | Mirjam Kurz |
| France | ALL | leaf/bud/cambium | 2019-2021 | Mirjam Kurz |
| Germany | GAI | bud | 2020 | Adrian Kölz |
| Germany | GRA | cambium | 2021 | Adrian Kölz |
| Germany | LEI | leaf, cambium | 2021 | Jonas Gorges |
| Germany | KIR | leaf | 2021 | Adrian Kölz |
| Germany | RIE | bud | 2020 | Adrian Kölz |
| Germany | STU | leaf | 2021 | Adrian Kölz |
| Germany | VOR | leaf, cambium | 2021 | Adrian Kölz |
| Germany | WUP | bud | 2020 | Jonas Gorges |

**Table S2.** Summary of the climatic conditions at the Oriental beech growing sites and the Western European plantations. Monthly mean values for seasons are represented by quarters of the year, i.e. the first quarter (i.e. January, February, March) as winter, the second as spring, the third as summer and the fourth as fall.

| Country | ID | Temperature (°C) | | | | | Precipitation (mm) | | | | |
|---|---|---|---|---|---|---|---|---|---|---|---|
| | | Winter mean | Spring mean | Summer mean | Fall mean | Annual mean | Winter mean | Spring mean | Summer mean | Fall mean | Annual Total |
| Iran | NOO | 7 | 18.9 | 24.8 | 12.9 | 15.9 | 71.4 | 48 | 54.2 | 105.8 | 838 |
| Georgia | LAG | -5.2 | 9 | 14.9 | 0.2 | 4.7 | 52.4 | 126.9 | 83.9 | 69 | 997 |
| Georgia | GOM | -2.3 | 11.8 | 17.5 | 3.1 | 7.5 | 33.5 | 88.9 | 68.1 | 39.7 | 691 |
| Russia | ING | 1 | 16.2 | 21.3 | 6.2 | 11.2 | 28.2 | 77.1 | 64.5 | 33 | 608 |
| Russia | LAN | 0.9 | 13.2 | 18.6 | 6 | 9.7 | 75.9 | 73.5 | 76.1 | 95.3 | 963 |
| Turkey | HAT | 2.7 | 14 | 20.7 | 7.6 | 11.2 | 139.5 | 81.7 | 31.3 | 109 | 1085 |
| Georgia | GOD | -5.9 | 7 | 12.9 | -0.2 | 3.4 | 92 | 101.9 | 89 | 105.9 | 1166 |
| Turkey | ORD | 1.5 | 12.3 | 18 | 7.1 | 9.7 | 107.3 | 82 | 72.9 | 137.9 | 1200 |
| Turkey | KAS | 0.7 | 11.8 | 17.2 | 5.7 | 8.9 | 90.8 | 58.7 | 64.2 | 124.6 | 1015 |
| Turkey | ZON | 4.9 | 15.8 | 21.3 | 10.1 | 13 | 94.7 | 63.2 | 66.5 | 113.5 | 1014 |
| Turkey | BUR | 2.9 | 13.5 | 19 | 7.9 | 10.8 | 113.7 | 82.5 | 42.9 | 113.2 | 1057 |
| Bulgaria | KOS | 5.3 | 16.5 | 22.4 | 10.6 | 13.7 | 44.1 | 39.7 | 27.2 | 62 | 519 |
| Bulgaria | NES | 1.8 | 13.8 | 19.4 | 7 | 10.5 | 51.3 | 60.3 | 43.3 | 68.3 | 670 |
| Switzerland | LEN | -1 | 9.1 | 13.3 | 2 | 5.8 | 120.9 | 128.3 | 131.7 | 126.9 | 1523 |
| Switzerland | WAE | 1.6 | 12.3 | 16.3 | 4.8 | 8.7 | 69.5 | 109.3 | 114.9 | 78 | 1115 |
| France | ALL | 2.8 | 13.1 | 16.9 | 5.5 | 9.6 | 54.2 | 65.6 | 65.7 | 57.9 | 730 |
| Germany | GAI | 1 | 12.1 | 15.9 | 4 | 8.2 | 72.9 | 84.7 | 87.1 | 76.3 | 963 |
| Germany | GRA | 1.9 | 12.8 | 16.6 | 4.9 | 9 | 53.5 | 98 | 97.1 | 57.9 | 920 |
| Germany | LEI | 3.4 | 13.9 | 17.7 | 6.2 | 10.3 | 44.7 | 82.3 | 85.4 | 52.8 | 796 |
| Germany | KIR | 2.8 | 13.1 | 17 | 5.6 | 9.6 | 55.2 | 52.7 | 59.1 | 61.5 | 686 |
| Germany | RIE | 2.7 | 13.4 | 17.1 | 5.6 | 9.7 | 48.6 | 64.6 | 72.2 | 57.7 | 729 |
| Germany | STU | 3.6 | 14.3 | 18.1 | 6.3 | 10.6 | 49.7 | 68.7 | 65.9 | 56 | 721 |
| Germany | VOR | 3.5 | 14.3 | 18 | 6.2 | 10.5 | 49.7 | 70.3 | 67.6 | 57.2 | 734 |
| Germany | WUP | 2.5 | 12 | 15.8 | 5.5 | 9 | 76 | 76 | 90.6 | 89.2 | 995 |

**Table S3.** Summary of the genetic diversity of 16 nuclear microsatellite loci in natural European beech (35 samples) and Oriental beech (103 samples) populations. k stands for the number of alleles; r stands for the allele size range.

| | **k** | | **r** | |
|---|---|---|---|---|
| **Locus** | **Oriental beech** | **European beech** | **Oriental beech** | **European beech** |
| csolfagus_29 | 9 | 3 | 16 | 8 |
| csolfagus_05 | 12 | 7 | 22 | 12 |
| csolfagus_06 | 16 | 7 | 34 | 16 |
| csolfagus_19 | 14 | 11 | 26 | 22 |
| csolfagus_31 | 13 | 10 | 32 | 22 |
| FS1_15 | 28 | 12 | 56 | 28 |
| sfc_0036 | 14 | 6 | 28 | 12 |
| sfc_1143 | 19 | 8 | 38 | 18 |
| DE576 | 9 | 5 | 24 | 18 |
| DUKCT | 9 | 6 | 16 | 20 |
| DZ447 | 24 | 5 | 56 | 6 |
| EEU75 | 14 | 7 | 32 | 18 |
| EJV8T | 10 | 7 | 18 | 12 |
| EMILY | 17 | 7 | 32 | 34 |
| ERHBI | 13 | 4 | 24 | 8 |
| concat14 | 9 | 4 | 18 | 22 |
| **MEAN** | 14.37 | 6.81 | 29.5 | 17.25 |

**Table S4.** Number of private alleles per genetic cluster and per locus, along with the frequency.

| Cluster | Locus | Allele | Frequency |
|---|---|---|---|
| Elbuz Mountains | csolfagus29 | 130 | 0.11 |
| Elbuz Mountains | csolfagus_06 | 229 | 0.07 |
| Elbuz Mountains | DE576 | 209 | 0.04 |
| Elbuz Mountains | DUKCT | 83 | 0.04 |
| Elbuz Mountains | EJV8T | 163 | 0.04 |
| Elbuz Mountains | EMILY | 166 | 0.04 |
| Elbuz Mountains | EMILY | 168 | 0.04 |
| Elbuz Mountains | ERHBI | 181 | 0.07 |
| Elbuz Mountains | sfc_0036 | 92 | 0.14 |
| Greater Caucasus | csolf_05 | 178 | 0.03 |
| Greater Caucasus | csolf_05 | 180 | 0.03 |
| Greater Caucasus | csolf_06 | 237 | 0.01 |
| Greater Caucasus | DE576 | 233 | 0.01 |
| Greater Caucasus | FS1_15 | 137 | 0.01 |
| Greater Caucasus | sfc_0036 | 90 | 0.01 |
| Greater Caucasus | sfc_0036 | 116 | 0.03 |
| Greater Caucasus | sfc_1143 | 137 | 0.01 |
| Taurus Mountains | concat14 | 193 | 0.04 |
| Taurus Mountains | DZ447 | 231 | 0.04 |
| Taurus Mountains | DZ447 | 241 | 0.04 |
| Taurus Mountains | DZ447 | 243 | 0.04 |
| Pontic Mountains | DZ447 | 211 | 0.02 |
| Pontic Mountains | ERHBI | 183 | 0.02 |
| Pontic Mountains | FS1_15 | 83 | 0.04 |
| Pontic Mountains | FS1_15 | 139 | 0.02 |
| Pontic Mountains | sfc_1143 | 147 | 0.02 |
| SW Black Sea coast | csolfagus29 | 150 | 0.02 |
| SW Black Sea coast | concat14 | 209 | 0.02 |
| SW Black Sea coast | concat14 | 211 | 0.02 |
| SW Black Sea coast | csolfagus_19 | 161 | 0.05 |
| SW Black Sea coast | csolfagus_19 | 187 | 0.02 |
| SW Black Sea coast | csolfagus_31 | 107 | 0.02 |

| | | | |
|---|---|---|---|
| SW Black Sea coast | csolfagus_31 | 111 | 0.04 |
| SW Black Sea coast | DZ447 | 187 | 0.02 |
| SW Black Sea coast | DZ447 | 225 | 0.09 |
| SW Black Sea coast | DZ447 | 233 | 0.02 |
| SW Black Sea coast | EEU75 | 101 | 0.18 |
| SW Black Sea coast | EMILY | 162 | 0.02 |
| SW Black Sea coast | ERHBI | 153 | 0.02 |
| SW Black Sea coast | FS1_15 | 127 | 0.02 |
| SW Black Sea coast | sfc_0036 | 118 | 0.04 |
| European beech | concat14 | 177 | 0.08 |
| European beech | csolfagus_31 | 121 | 0.27 |
| European beech | csolfagus_31 | 131 | 0.02 |
| European beech | DUKCT | 77 | 0.04 |
| European beech | DZ447 | 190 | 0.06 |
| European beech | DZ448 | 204 | 0.02 |

**Table S5.** Analyses of molecular variance.

| Source of variation | d.f. | Sum of squares | Variance components | | Percentage of variation |
|---|---|---|---|---|---|
| **Among populations** | 5 | 229.42 | 0.88 | Va | 13.78 |
| **Among individuals within populations** | 133 | 777.00 | 0.33 | Vb | 5.09 |
| **Within individuals** | 139 | 721.50 | 5.19 | Vc | 81.13 |
| **Total** | 277 | 1727.92 | 6.40 | | |

**Fixation Indices**

$F_{IS}$: 0.0591

$F_{ST}$: 0.1378

$F_{IT}$: 0.1887

Fig. S1. Sampling scheme. We sampled seedlings around the largest adult trees (focal mother trees) and along four transects facing four different directions around it. We sampled seedlings within circular plots, each with a radius of 2 m, whose centers were located 2, 6 and 14 m away from the focal tree. The circular plots were labelled A–L. We started with the A-B-C transect and then continued in a clockwise direction. The starting orientation of the first transects was randomly chosen.

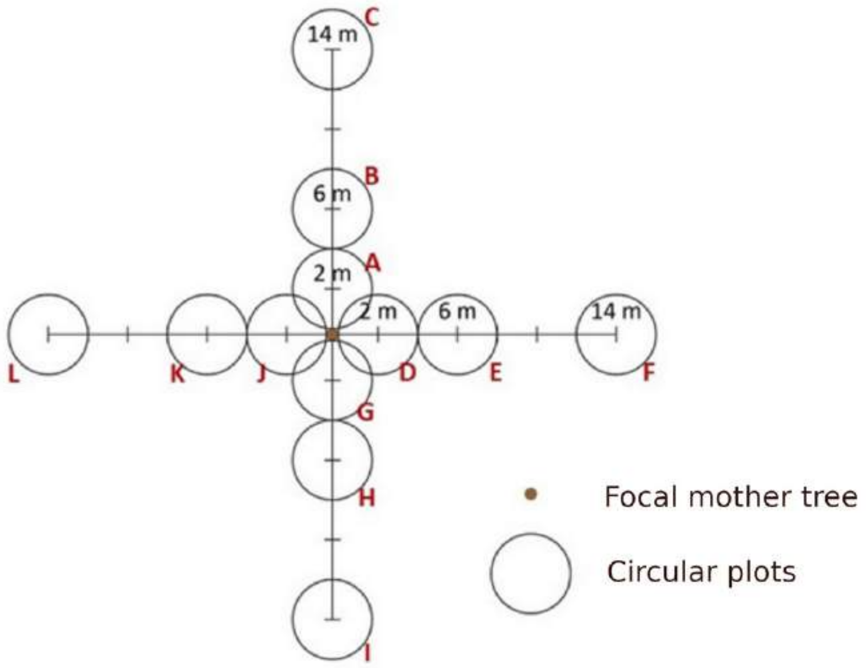

Fig. S2. Definitions of Fagus phenological stages following Vitasse et al. (2013). Stage 1: bud swelling, Stage 2: bud burst, Stage 3: leaf-out, Stage 4: leaf unfolded. In seedlings, we pressed on the bud, if it was soft we noted stage 1, if it was hard we noted stage 0.

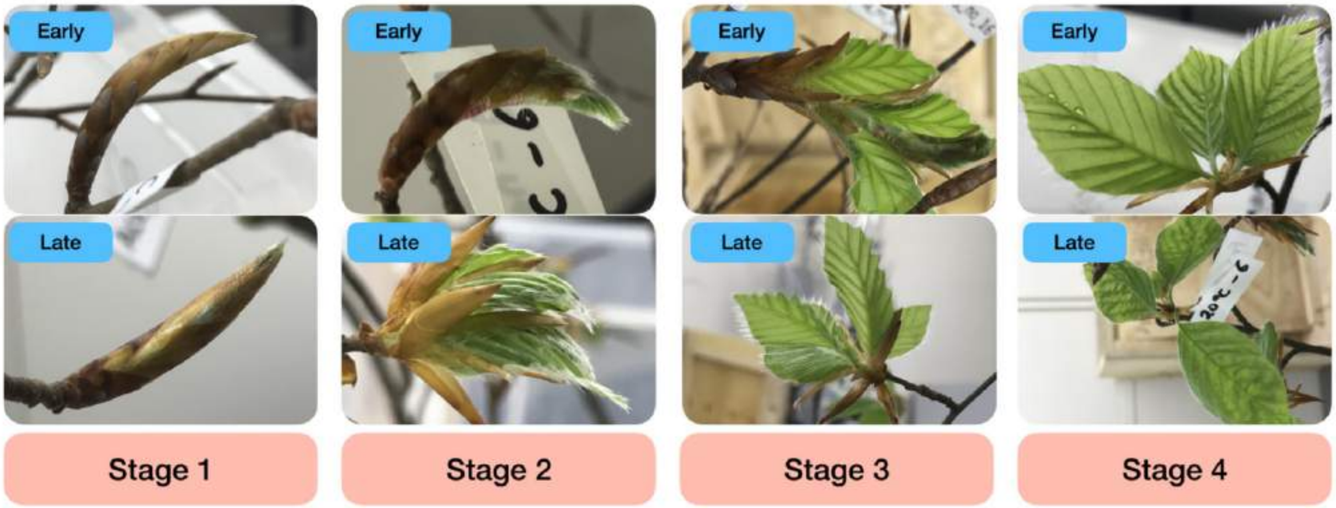

Fig. S3. Allele frequency distributions per cluster. See Fig. 1 for the geographic range of the clusters.

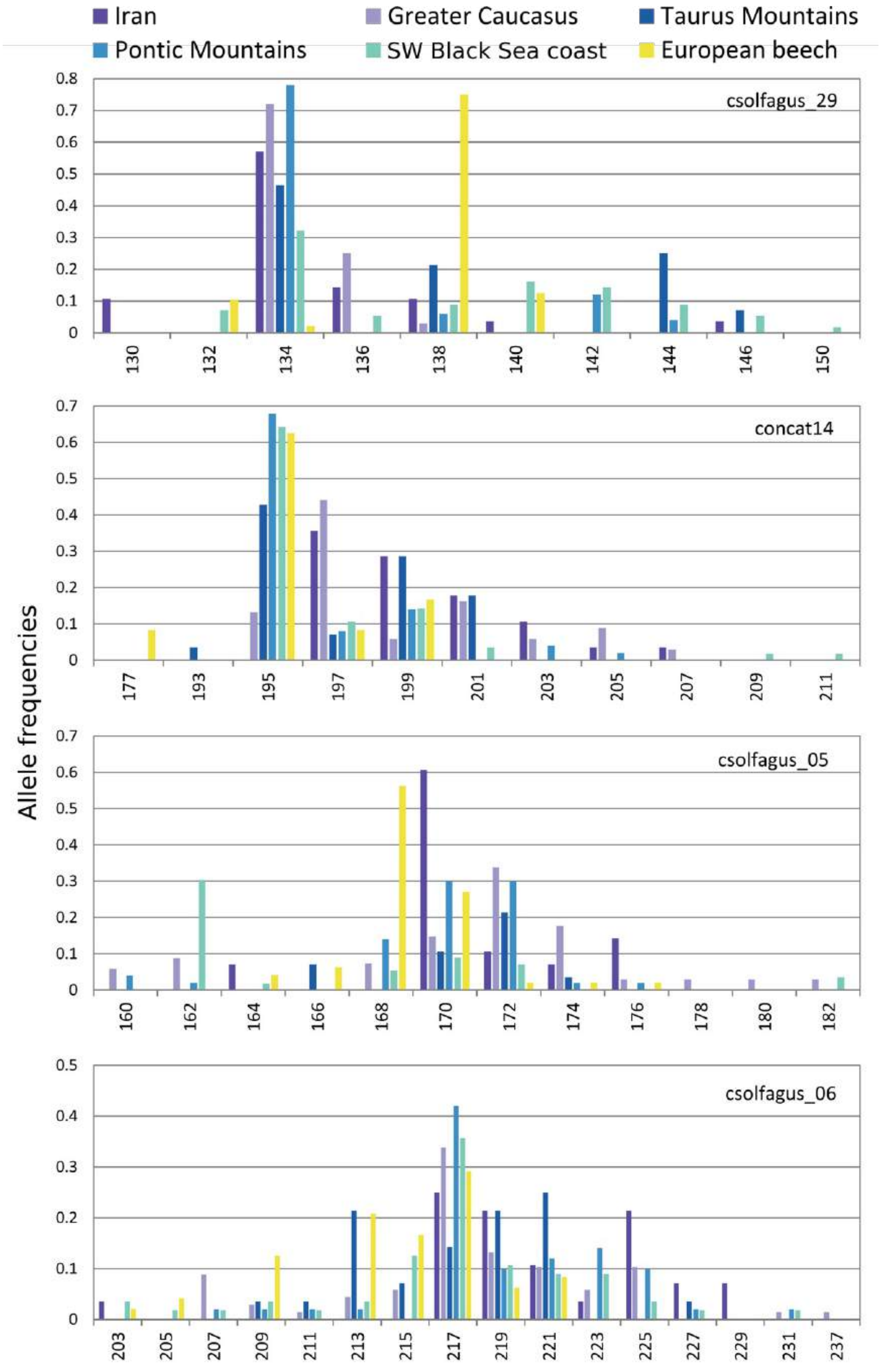

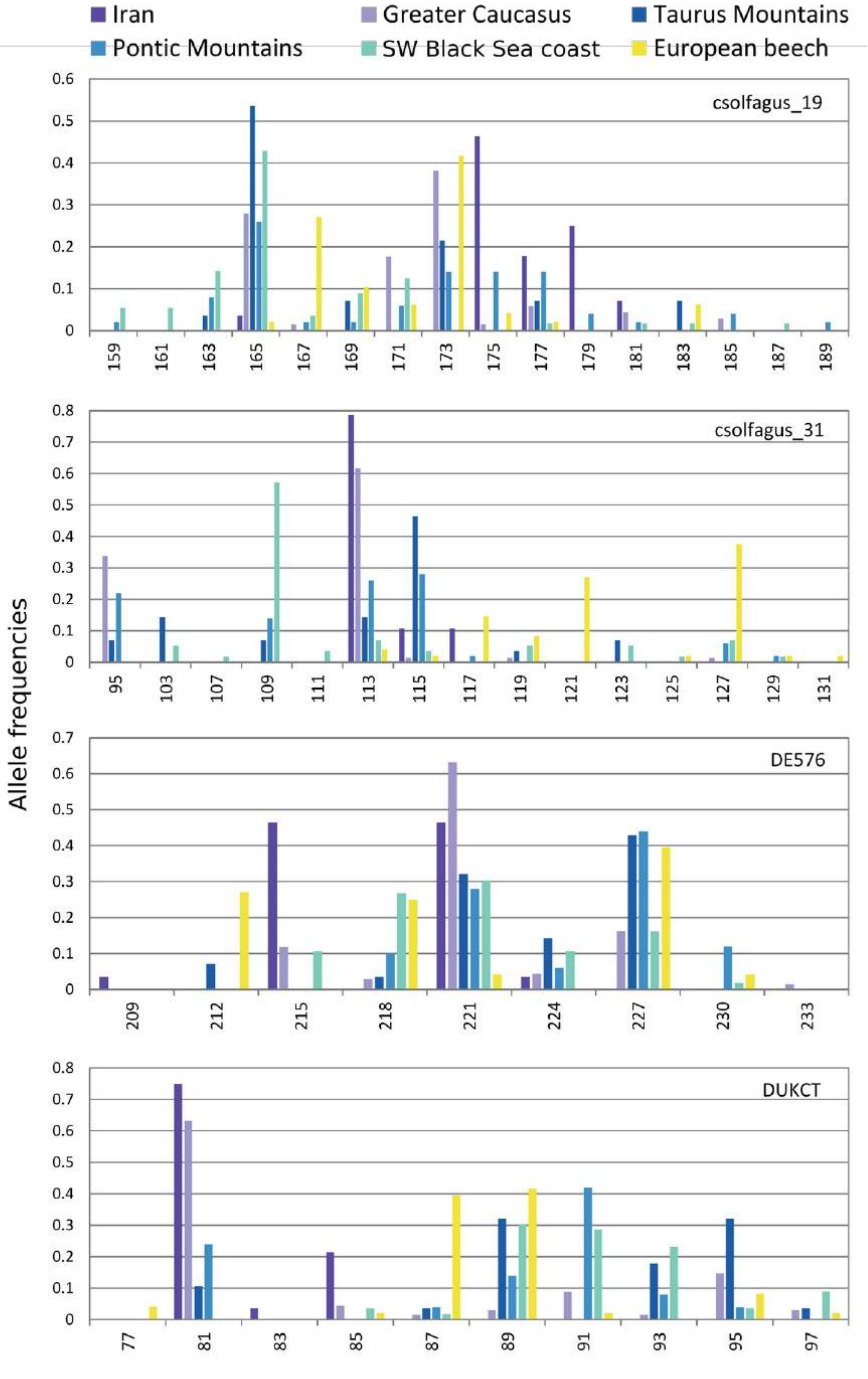
Iran
Greater Caucasus
Taurus Mountains
Pontic Mountains
SW Black Sea coast
European beech
Allele frequencies
csolfagus_19
csolfagus_31
DE576
DUKCT

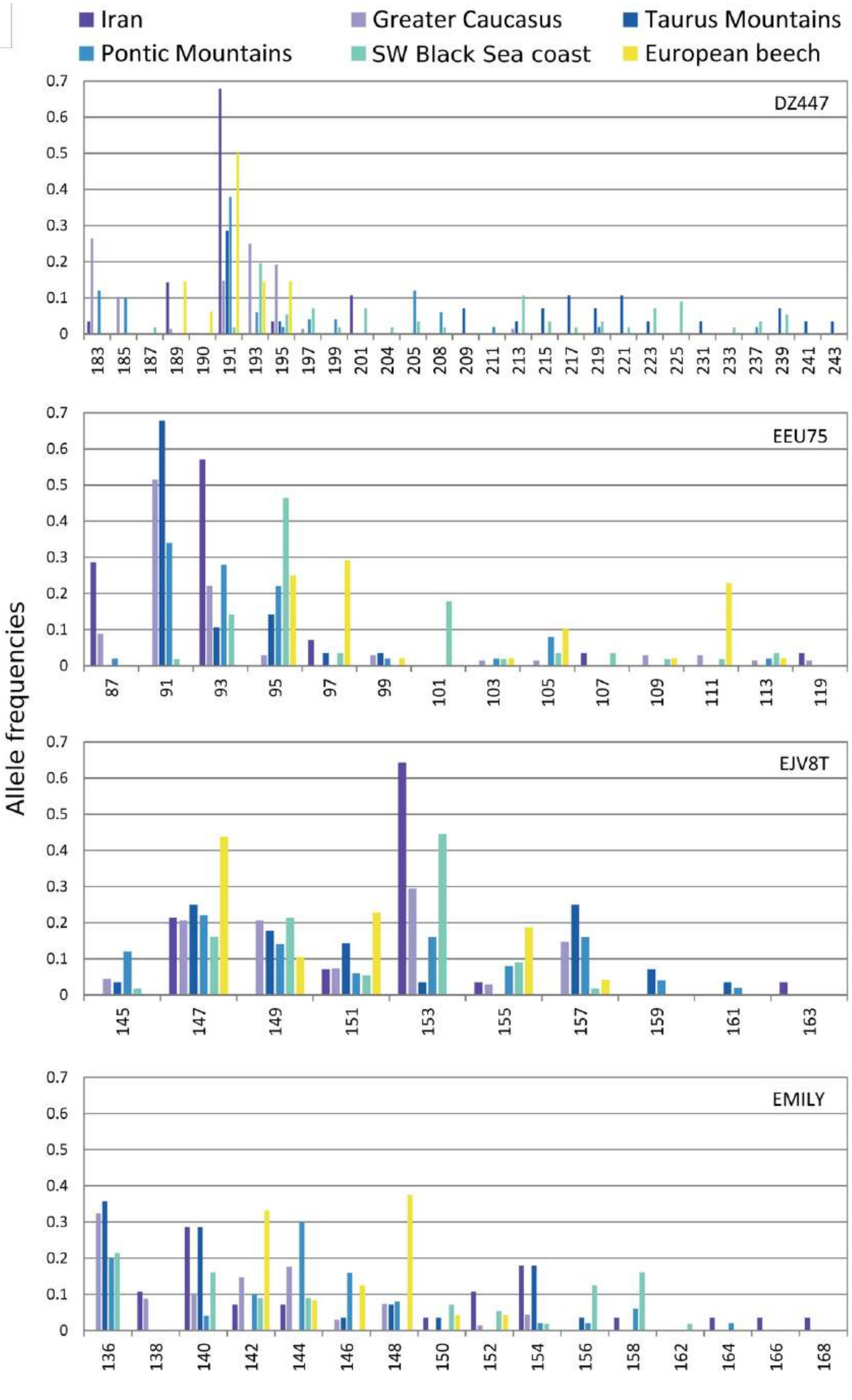
Iran
Greater Caucasus
Taurus Mountains
Pontic Mountains
SW Black Sea coast
European beech
Allele frequencies
DZ447
EEU75
EJV8T
EMILY

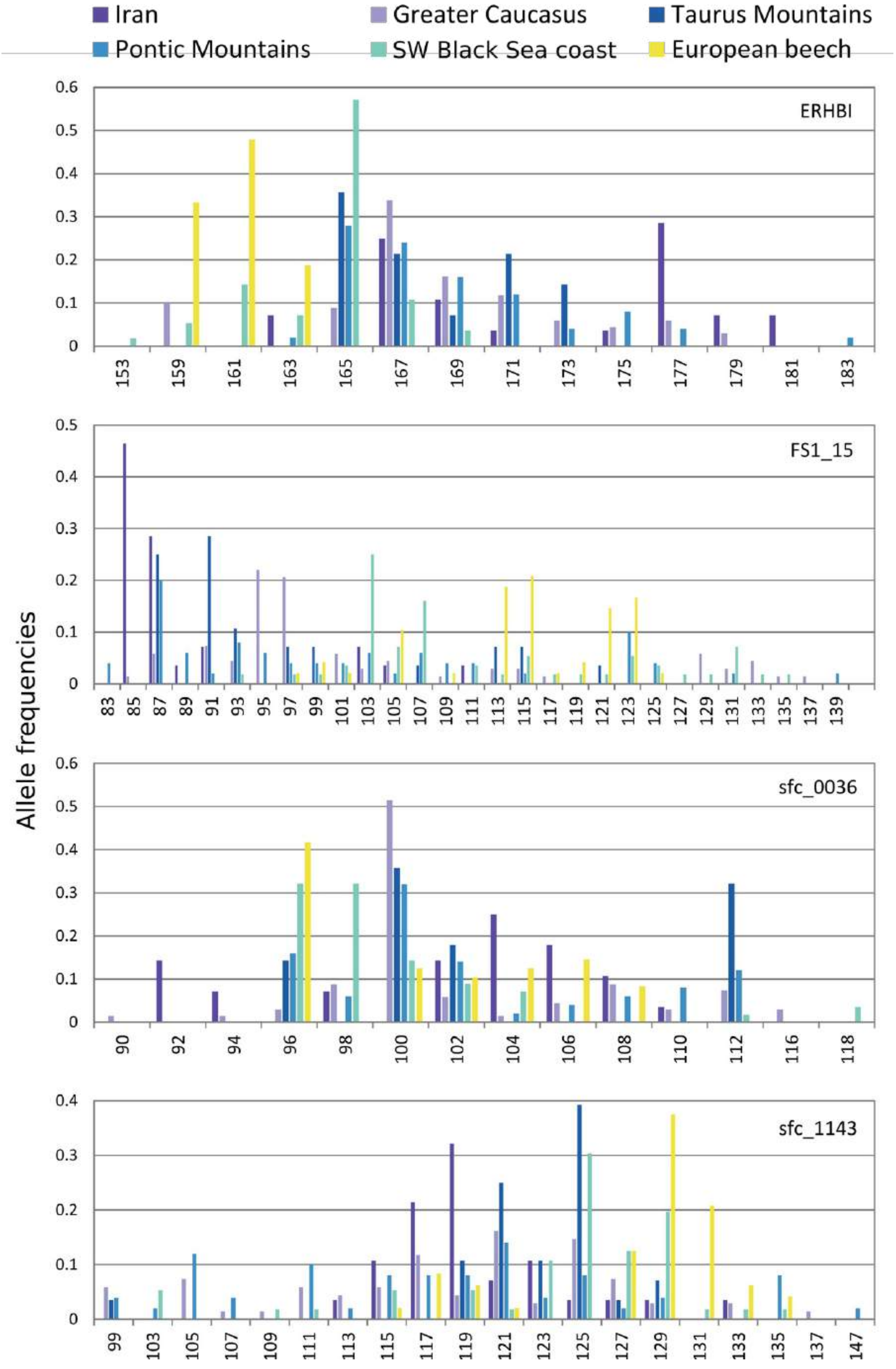

Iran
Greater Caucasus
Taurus Mountains
Pontic Mountains
SW Black Sea coast
European beech
ERHBI
FS1_15
sfc_0036
sfc_1143
Allele frequencies
153 159 161 163 165 167 169 171 173 175 177 179 181 183
83 85 87 89 91 93 95 97 99 101 103 105 107 109 111 113 115 117 119 121 123 125 127 129 131 133 135 137 139
90 92 94 96 98 100 102 104 106 108 110 112 116 118
99 103 105 107 109 111 113 115 117 119 121 123 125 127 129 131 133 135 137 147

Fig. S4. Genetic clustering using the software Structure and settings identical to the analysis shown on Fig. 1B, but using K from 2 to 6.

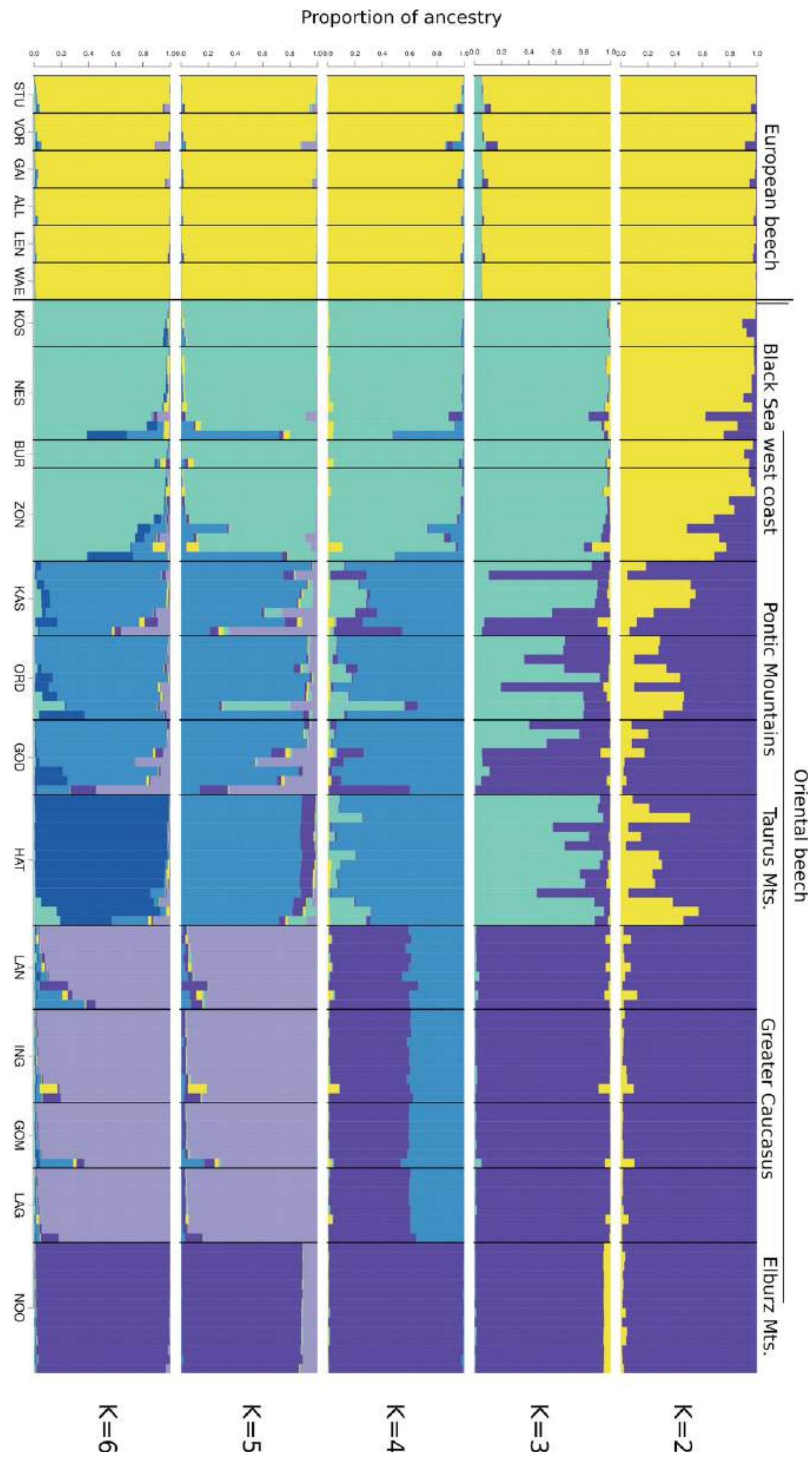

Fig. S5. Summary statistics for the performance of the genetic clustering using the software Structure for the analyses shown on Fig. 1b.

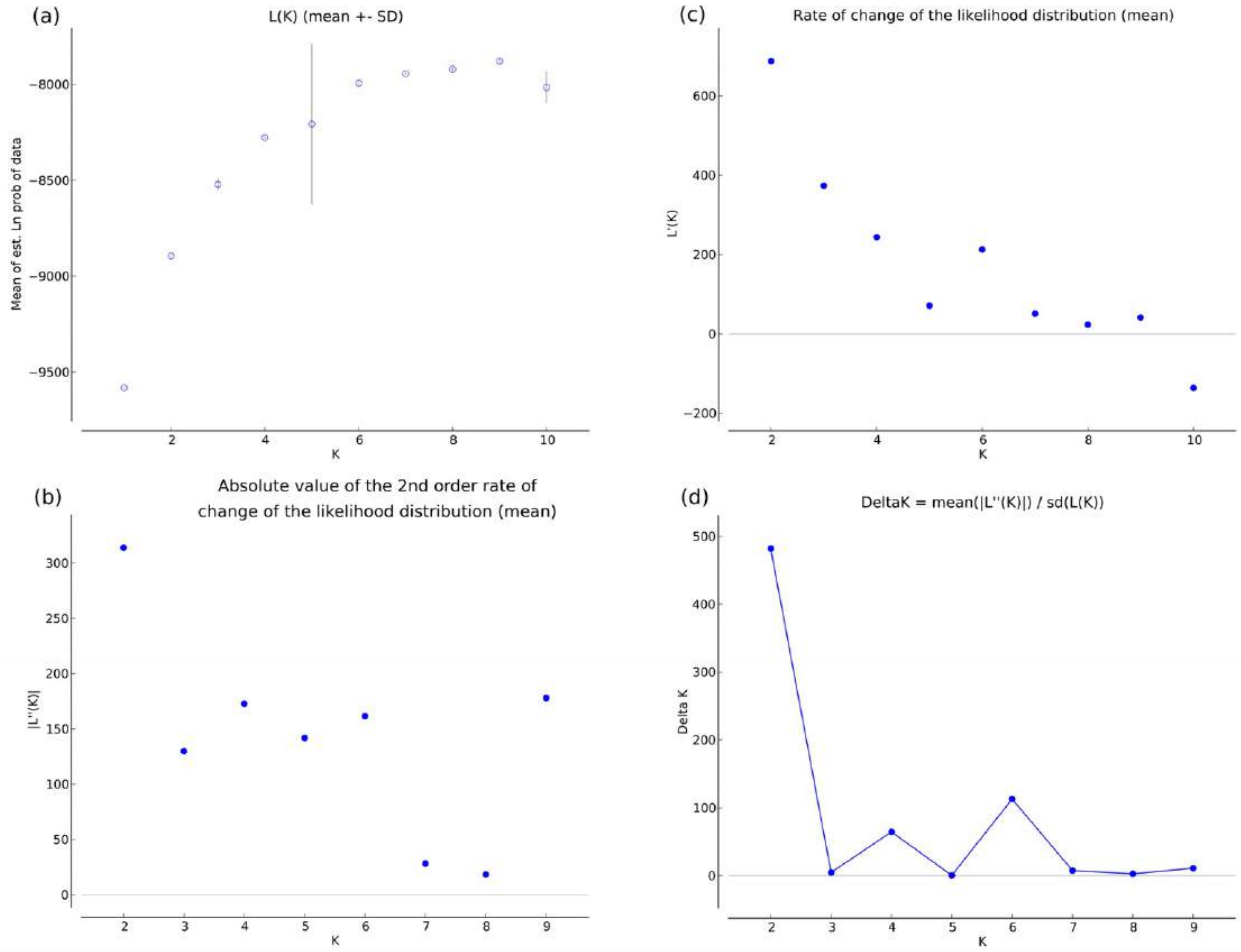


(e) Evanno's Table

| K | Reps | Mean LnP(K) | Stdev LnP(K) | Ln'(K) | \|Ln''(K)\| | Delta K |
|---|---|---|---|---|---|---|
| **1** | 20 | -9582.455 | 1.0298 | | | |
| **2** | 20 | -8894.875 | 0.6512 | 687.58 | 313.95 | 482.099999 |
| **3** | 20 | -8521.245 | 28.0399 | 373.63 | 129.91 | 4.633037 |
| **4** | 20 | -8277.525 | 2.6819 | 243.72 | 172.83 | 64.443511 |
| **5** | 20 | -8206.635 | 417.1175 | 70.89 | 141.775 | 0.339892 |
| **6** | **20** | **-7993.97** | **1.4313** | **212.665** | **161.49** | **112.830071** |
| **7** | 20 | -7942.795 | 3.7286 | 51.175 | 28.13 | 7.544347 |
| **8** | 20 | -7919.75 | 6.6464 | 23.045 | 18.225 | 2.742066 |
| **9** | 20 | -7878.48 | 16.3444 | 41.27 | 177.965 | 10.888414 |
| **10** | 20 | -8015.175 | 79.221 | -136.695 | | |

Pr

Fig. S6. Origin of Oriental beech in 11 Western European plantations. Genetic clustering was performed using the software Structure, as shown on Fig. 2, but with USEPOPINFO.

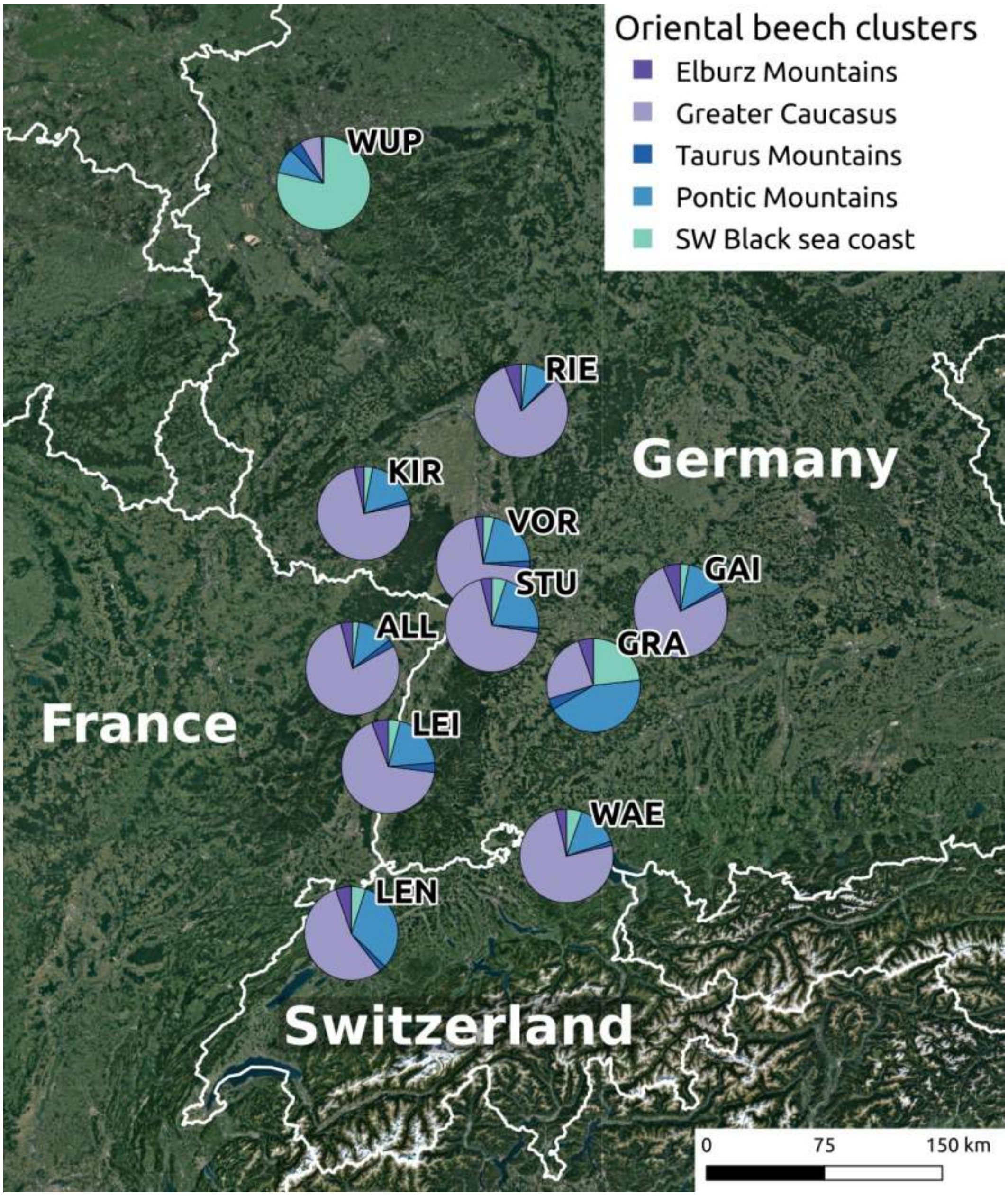

Fig. S7. Ancestry coefficients from the software Structure for offspring sampled around the focal mother trees in WAE and ALL. See Fig. 3 for an overview of the results.

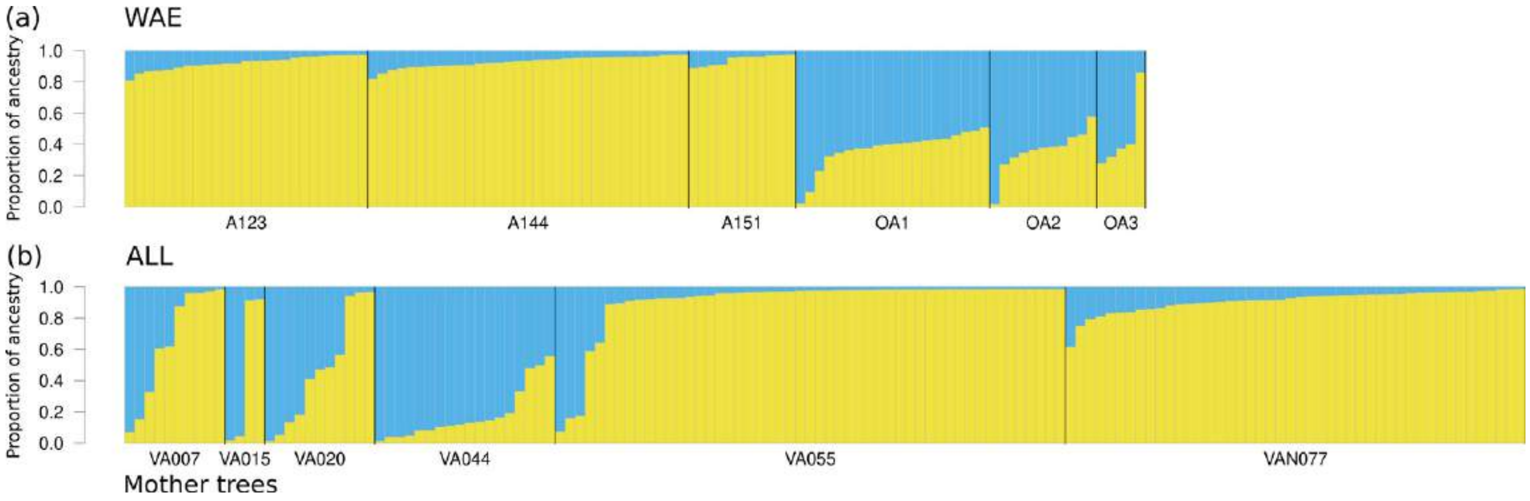
